# Reduced-order modeling of solute transport within physiologically realistic solid tumor microenvironment

**DOI:** 10.1101/2025.11.15.688588

**Authors:** Mohammad Mehedi Hasan Akash, Mohammad Yeasin, Shima Mahmoudirad, Redowan A Niloy, Jiyan Mohammad, Katie Reindl, Anupum Pandey, Saikat Basu

**Author notes:** Corresponding author; Web:https://sites.google.com/view/basulab-tour/home.

## Abstract

Solid tumors are characterized by densely packed extracellular matrices and limited vascularization, creating significant resistance to both diffusive and convective transport. In this study, we developed an integration of numerical computations with a theoretical modeling framework that couples three phase viscous-laminar transient simulations of glycocalyx-patched tumor vessel resolving plasma, red blood cells (RBCs), and white blood cells (WBCs) and tracking their volume fractions to a calibrated reverse advection-diffusion (RAD) model for intratumoral plasma transport. The reduced-order tumor microenvironment model uses histology-informed extracellular matrix (ECM) tumor domain and packing fraction, together with explicit glycocalyx-patch electrohydrodynamics (EHD) at the tumor vessel wall. At the fenestra, EHD increases inlet plasma intensity relative to a non-EHD framework across all models (means: 0.576 non-EHD vs 0.722 EHD; gain 25.34%). Numerical simulations of plasma perfusion in both the tumor ECM domain and a microfluidic benchmark exhibit two-stage kinetics, with an initial advection-dominated regime. The RAD model reproduces this behavior and, after a simple temporal calibration to account for pore-scale hydrodynamic acceleration resolved by computational fluid dynamics (CFD), matches the observed propagation. By using fully resolved, EHD-inclusive multiphase CFD simulations to calibrate a reduced-order RAD model parameterized by measurable geometric features, we bridge the gap between classical Darcy–Starling tissue perfusion models and fully resolved CFD. The resulting framework provides a tractable, mechanism-grounded tool for quantifying plasma progression in dense solid tumors.

## 1 Introduction

Solute transport in solid tumors is fundamentally governed by the structural and mechanical properties of the surrounding tissue [Gullino, P. M., 1980, Sefidgar et al., 2014]. Irregular vascular architecture, dense stromal packing, and elevated interstitial resistance disrupt flow pathways and reduce transport efficiency [Vaupel, P.and Kallinowski, F. and Okunieff, P., 1990, Jain et al., 2014, Koumoutsakos, P. et al., 2013, Stylianopoulos et al., 2018a, Jiang et al., 2020]. These conditions create spatial variability in pressure-driven and diffusive mechanisms, producing highly non-uniform intratumoral distributions. Despite the enhanced permeability and retention effect often reported for nanoparticle delivery [Maeda, 2012], perfusion in tumors can be much lower than in surrounding healthy tissues due to vascular constriction and abnormal leakiness into the extracellular space [Stylianopoulos, T. and Jain, R. K., 2013].

Solid tumors, which constitute the majority of cases, present as noncystic masses categorized as benign or malignant [Gavhane, Y. N. et al., 2011, Rahman et al., 2023] and exhibit high resistance to diffusion [Jain, R. K., 1996, 1998] with limited blood and lymphatic drainage that impedes advection [Chim and Mikos, 2018]. Accurate, mechanics-based quantification of spatiotemporal solute distributions is therefore essential for clinical decision-making [d’Esposito et al., 2018], as flow–structure interactions and vascular leakiness elevate interstitial pressure and vary across patient-specific tumor topography [Soltani and Chen, 2011, Zhang et al., 2013]. Beyond hydrodynamics, electrohydrodynamics (EHD) introduces electrical body forces that can bias plasma transport: in vivo, the endothelial glycocalyx presents patched and heterogeneous surface charge that can drive EHD responses at the vessel wall [Weinbaum et al., 2007, Huilgo, R.R. and Phan-Thien, N., 1997, Sumets et al., 2018, Teodoro et al., 2024]. Blood flow is multiphase and non-Newtonian: plasma with suspended RBCs and WBCs creates a heterogeneous mixture in which collisions and aggregation change bulk rheology and dispersion [Merrill et al., 1963, Baskurt, O. K. and Meiselman, H. J., 2003]. Two phase hemodynamics models [Jung et al., 2006] improve over single-phase assumptions yet remain incomplete; three-phase formulations [Akash et al., 2023] better capture particulate interactions and motivate coupling geometry and biology in a predictive framework. Prior multiphase–EHD studies [Rahmat, Sabanci University, 2017] provide valuable numerical background but typically do not incorporate explicit three-phase blood or patched glycocalyx charge [Kang et al., 2018]. In parallel, classical mean-transport models (Darcy’s law [Baish et al., 2011]; Starling’s equation [Sugihara-Seki and Fu, 2005]) remain useful at the tissue scale but overlook key features of tumor hemodynamics, including non-Newtonian, particulate rheology, and geometry-specific microstructure. Likewise, single-phase computational fluid dynamics (CFD), which treats blood as a homogeneous continuum, cannot resolve the interactive dynamics and spatial distribution of suspended particulate matter [Womersley, J. R., 1957, Attinger, E. O., 1964, Srivastava, V. P., 2007].

Guided by these limitations, we develop a three-phase simulation framework that explicitly resolves blood mechanics and glycocalyx-mediated EHD inside tumor vessels to mechanistically quantify—and, under controlled conditions, enhance plasma ingress into the tumor extracellular matrix (ECM) in a biologically faithful. Building on these high-fidelity simulations, we introduce a CFD-informed reverse advection-diffusion (RAD) model: an advection–diffusion framework whose boundary data and closures are calibrated from the resolved vessel physics and whose coefficients are expressed in terms of measurable geometric descriptors such as stromal packing fraction [Sefidgar et al., 2014, 2015, Soltani and Chen, 2011]. Here, “reverse” denotes using vessel-scale physics to inform tissue-scale transport closures. Our aim is to retain the mechanistic fidelity of vessel-resolved three-phase CFD—including localized EHD at patched glycocalyx segments–while bridging the gap between classical Darcy–Starling formulations and modern fluid dynamics by embedding these vessel-scale effects into a reduced-order transport model. This motivates the following questions:

Q1. As a marker of enhanced biological realism, how would the inclusion of electrohydrodynamic (EHD) effects change the perfusion trend at the fenestra, compared to the simpler non-EHD framework?

Q2. As a parametric tool, could we set up a simplified, reduced-order convection-diffusion model for plasma transport in the ECM, with inlet flow conditions guided by the EHD-inclusive CFD simulations?

To address these questions, we first perform three phase CFD of tumor-vessel flow with EHD localized to the vessel domain to quantify changes in plasma-perfusion metrics relative to a non-EHD baseline; we then develop and calibrate the RAD model to express penetration and advection diffusion transitions explicitly in terms of packing fraction and geometry, thereby bridging the gap between classical mean-transport descriptions and fully resolved CFD. Finally, we benchmark the model’s key predictions (front advance, wetted-area scaling, and transition time) against complementary microfluidic validation. Preliminary results from this work have been presented at the Annual Meetings of the American Physical Society’s Division of Fluid Dynamics [Akash et al., 2024, Akash, M. M. H. et al., 2021].

## 2 Methods

### 2.1 Domain reconstruction

#### 2.1.1 Development of reduced-order tumor vessel

To interrogate electrohydrodynamics (EHD)–driven changes in plasma perfusion, we employed a biomimetic tumor microvessel coupled to an extracellular matrix (ECM) domain (Fig. 1C). The reduced-order vessel geometry follows the configuration in our prior work and its validated *in silico* capillary model [Akash et al., 2023, 2024]: a 500 *µ*m-long channel with a single fenestra—a rounded wall depression—that serves as the sole plasma–ECM exchange interface. The fenestra is 2.6 *µ*m deep and 5.2 *µ*m in diameter, sized to permit plasma entering while excluding larger formed elements such as red blood cells (RBCs, 7–8 *µ*m) [Nithiarasu, 2022]. Endothelial-scale context (endothelial cells, glycocalyx, RBCs, and white blood cells (WBCs)) is illustrated in Fig. 1A, and the near-fenestra mesh is shown in Fig. 1D. To represent heterogeneous endothelial surface properties that mediate EHD, we applied wall-attached glycocalyx patches with fixed streamwise extent of 0.5 *µ*m Dull and Hahn [2022], Weinbaum et al. [2021]. Patches were distributed in a randomized, nonperiodic pattern (some adjacent, others separated by a single gap) and held fixed across all cases for reproducibility. Numerically, patched regions were implemented via a user-defined forcing term localized to wall-adjacent control volumes, adding an electric body force consistent with the prescribed patched surface charge (details in Sec. 2.2.1). In the schematic and layout, red wall segments denote patched, glycocalyx-bearing regions where EHD forcing is applied (Fig. 1A,C), whereas black wall segments in Fig. 1C indicate non-patched wall with no EHD forcing.

**Figure 1:**
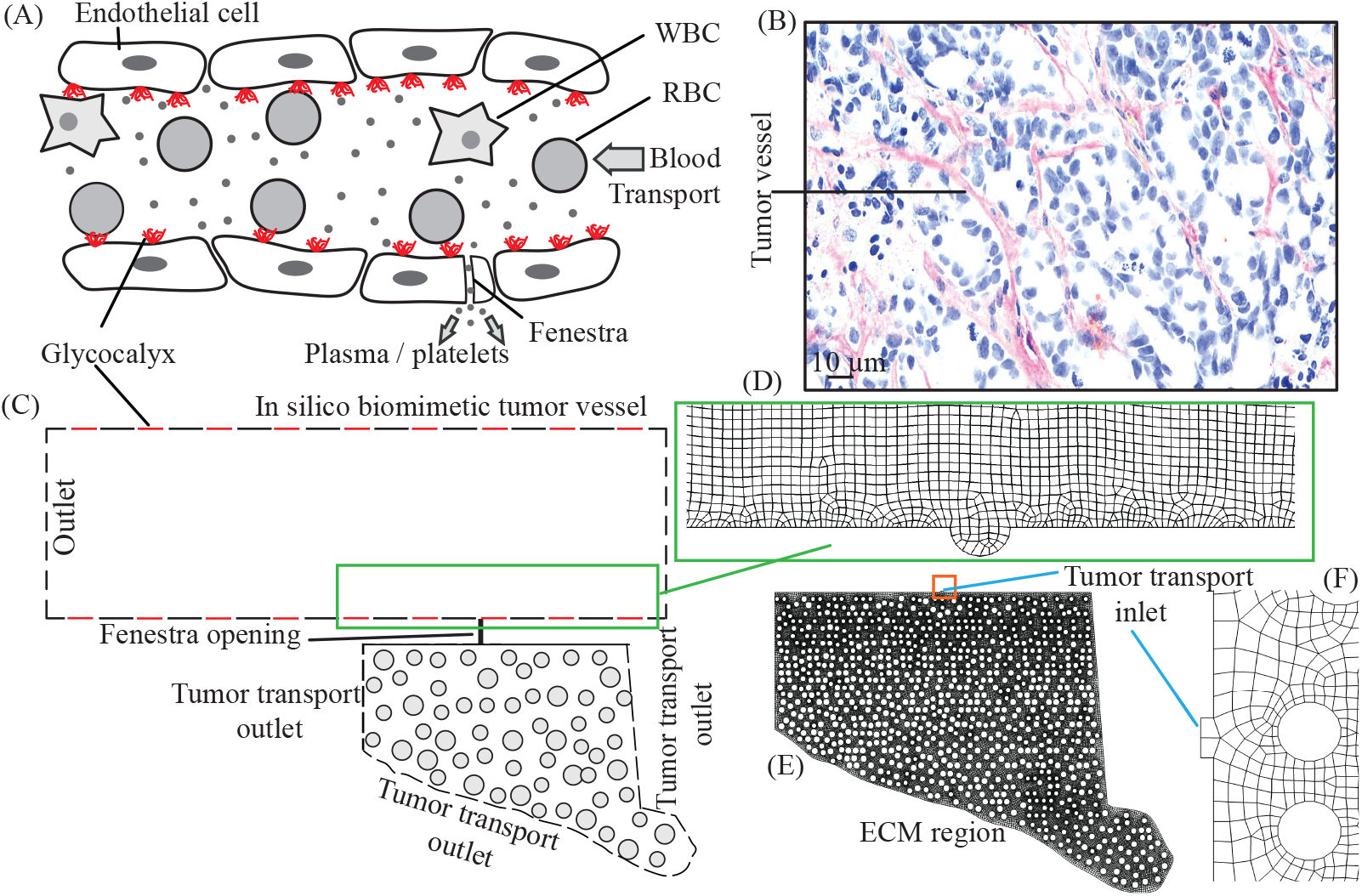
Physiologically inspired schematic and simulation geometry of a tumor vessel coupled to the surrounding tumor ECM domain: (A) Endothelial capillary schematic showing the endothelial glycocalyx (red patches), red blood cells (RBCs), white blood cells (WBCs), and a plasma-only fenestra that connects the vessel to the tumor extracellular matrix (ECM). This schematic motivates the reduced-order model; in panel (C), red lines indicate glycocalyx-patched (electrohydrodynamic) wall segments, and black lines indicate non-patched segments. (B) Representative histology of a tumor vessel and a scanned cross-section of a dense pancreatic tumor implanted in mice; this morphology informed the simulated vessel. (C) Two dimensional biomimetic domain of the tumor microenvironment, where the reduced-order tumor vessel is connected to the tumor stroma via a fenestra. For numerical simulations, the domain in (C) is partitioned into two segments. The modeling strategy restricts the entry of suspended particulates at the fenestra by selecting a fenestra diameter smaller than typical particle sizes. (D–F) The two segments of the test geometry with their corresponding meshes: (D) a portion of the tumor vessel showing the circular wall depression that coincides with the fenestra location; (E) the ECM region with the fenestra (tumor-transport inlet); and (F) a zoomed-in view of the fenestra region with finer mesh elements. Additional details of the ECM geometry are provided in Sec. 2.1.2.

Sensitivity of perfusion to the location of plasma entry was probed by generating *N* = 15 geometries in which the fenestra center was shifted axially along the vessel while keeping fenestra shape and the patch pattern unchanged (Fig. 2A). With 25 *µ*m clearance zones at inlet and outlet, admissible centers span [25, 475] *µ*m. We define

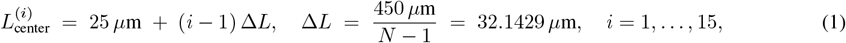

which places one nonoverlapping fenestra per configuration over the effective length *L*_eff_ = 450 *µ*m. The first admissible center at 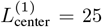 *µ*m is Model 1, and the sequence proceeds in steps of Δ*L* = 32.1429 *µ*m so that Model 15 corresponds to 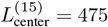 *µ*m.

**Figure 2:**
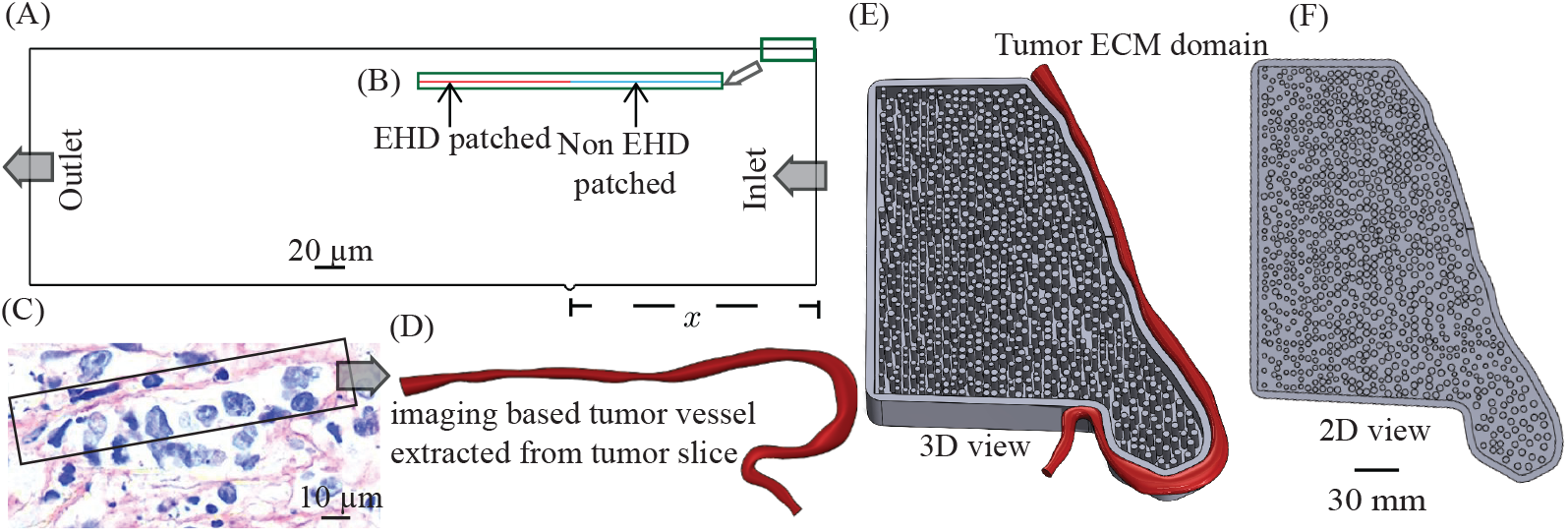
Physiologically inspired tumor vessel and ECM geometries linking histology to reduced-order models: (A) Reduced-order tumor vessel with inlet (right) and outlet (left). The bracketed axis *x* marks the admissible interval for the fenestra center used in the parametric sweep; varying *x* changes the model index (Models 1–15 per Eq.1). (B) Zoomed view from the actual CAD model of the luminal wall showing alternating surface segments: red line shows EHD-patched, glycocalyx-bearing regions where electrohydrodynamic forcing is applied; blue line shows non-EHD patched wall with no EHD forcing (implementation details in Sec. 2.2.1). (C) H&E histology of an orthotopic human pancreatic tumor slice illustrating irregular microvasculature; this specimen informed the vessel geometry. (D) Imaging-based vessel extracted from the tumor slice and used to define the curved vessel–ECM interface. (E) Tumor ECM domain (3D view) populated with circular fiber bundles to achieve the prescribed packing fraction; the curved red vessel indicates the vessel–ECM boundary used to locate the fenestra. (F) Planar (2D) surrogate of the ECM domain used for transport simulations; the curved red vessel is not adapted as a cross-section here because the 2D surrogate contains only the ECM region while retaining the same boundary imposed at the fenestra (Sec. 2.1.2).

All geometries were constructed and meshed in ANSYS Workbench 2023 R1 using DesignModeler (DM) for CAD and the integrated mesher (ANSYS Inc., Canonsburg, Pennsylvania). Because the computational domain is two-dimensional (Fig. 1C–F), unstructured triangular meshes were employed, with local refinement around the fenestra and patched-wall regions to resolve steep pressure/velocity gradients while keeping the far field economical. A systematic mesh-refinement study (Table 1) established a numerically converged resolution.

**Table 1:** Mesh-refinement study: domain-averaged pressure and velocity. The 1.00 *µ*m mesh (~ 77k elements) is selected; differences to the 0.76 *µ*m mesh are negligible.

| Element Size ($\mu\text{m}$) | Total Elements | $P_{\text{avg}}$ (Pa) | $V_{\text{avg}}$ (m/s) |
| --- | --- | --- | --- |
| 5.0 | 8k | 2193.67 | $3.1 \times 10^{-5}$ |
| 3.0 | 19k | 2180.66 | $4.3 \times 10^{-5}$ |
| 1.00 | 77k | 2180.26 | $3.43 \times 10^{-5}$ |
| 0.76 | 80k | 2180.48 | $3.44 \times 10^{-5}$ |

Between 1.00 *µ*m and 0.76 *µ*m, the change in domain-averaged pressure is 0.22 Pa (about 0.010% of 2180 Pa), and the change in average velocity magnitude is 1.0 × 10^−7^ m/s (about 0.29% relative to 3.43 × 10^−5^ m/s). Given these sub-percent differences and the ~ 4% increase in element count for 0.76 *µ*m, the 1.00 *µ*m mesh (77k elements) was adopted for all 15 geometries as the best trade-off between accuracy and computational cost, ensuring uniform resolution across the parametric sweep.

#### 2.1.2 Development of the tumor extracellular matrix (ECM) domain

To construct the ECM domain, we first captured the specimen-specific vessel shape. We developed a slice–to–geometry pipeline (Fig. 2C → 2D) to reproduce the vessel boundary and to build the ECM domain used in transport simulations (Fig. 2E shows a 3D view). The slice (Fig.2C) derives from human pancreatic cancer cells orthotopically injected into the pancreas of nude mice. Tissue was collected, fixed in formaldehyde for 24 h, paraffin embedded, and sectioned at 5 *µ*m. Sections were deparaffinized in Histo^®^ Clear and ethanol, rehydrated, stained with hematoxylin (5 min) and eosin (1 min), dehydrated in 95–100% ethanol, cleared in xylene, mounted, and imaged on a Zeiss Axio Observer Z1 microscope. Histology preparation, staining, and imaging follow our prior work [Akash et al., 2023].

An in-house framework detected the luminal periphery using intensity-based edge finding with morphological cleanup. The resulting edge contours were converted to ordered coordinates along the vessel wall and imported into CAD, where they were aligned to a common reference frame. An interpolation step then produced a continuous spline curve with *C*^1^ continuity that faithfully represented the vessel geometry while removing pixelation artifacts from the raster image. This curve defined the top, curved boundary of the ECM domain used in the numerical simulations. Using this boundary as the vessel–ECM interface, we constructed the adjacent ECM region and populated it to achieve a prescribed in-plane packing fraction. Stromal crowding was represented by *N*_fib_ = 870 circular bundles placed within the ECM such that

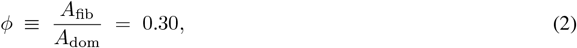

with a target mean bundle diameter 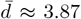 *µ*m (range 3.0–4.5 *µ*m). Here *A*_fib_ is the total fiber cross-sectional area and *A*_dom_ is the ECM domain area. Fixing *ϕ* sets the porosity *ε* = 1 − *ϕ*, which enters the transport closures for the effective diffusivity *D*_eff_ (via tortuosity) and the hydraulic permeability *k*, thereby limiting both diffusive penetration and pressure-driven perfusion. The imposed fiber field was checked against orientation statistics from the same segmentation workflow and against reported stromal-alignment trends in tumor tissues [Eekhoff, J. D. and Lake, S. P., 2020]. Coupling to the vessel is applied along the extracted boundary locus at the fenestra by prescribing the plasma-field inlet (and associated flux) from the vessel simulation; this enforces a smooth interface without explicitly meshing the vessel solid within the ECM grid. The two dimensional ECM domain (Fig. 2F) was meshed with approximately 5.9 × 10^4^ unstructured triangular elements (see also Fig. 1E), with local refinement near the vessel–ECM interface. Near the boundary locus corresponding to the fenestra, a characteristic length scale *a*_fen_ = 0.1 *µ*m was used in the near-wall mass-transfer closure and scaled locally with the extracted wall geometry. In the simulations, the multiphase transport calculation in the vessel yields the plasma volume fraction at the artificial circular depression (fenestra), which is then used as the inlet condition for the subsequent ECM computation (Fig. 1E), where intratumoral plasma transport is tracked.

### 2.2 Simulation setup

#### 2.2.1 Simulation method for blood flow inside the tumor vessel

We model the in silico transport of blood within tumor vessels as an unsteady, viscous–laminar, two dimensional (planar) flow. Numerical solutions are obtained in ANSYS FLUENT 2023 R1 using its Eulerian multiphase model (no custom inter-particle equations). Plasma is the primary (continuous) phase; red blood cells (RBCs) and white blood cells (WBCs) are treated as two dispersed phases. The formulation follows the classical multiphase CFD framework [Gidaspow, D., 1994, Anderson, T. B. and Jackson, R., 1967] and is appropriate for in vivo hematocrits of 30–55% [Jung et al., 2006]. Closures (mixture shear-thinning, interphase drag, agglomeration shape factor) follow [Akash et al., 2023].

Let *k* ∈ {*p, r, w*} index plasma (*p*), RBCs (*r*), and WBCs (*w*), with volume fractions *α*_*k*_ and velocities **u**_*k*_ = (*u*_*k*_, *v*_*k*_) in 2D, and let ∇ = (*∂*_*x*_, *∂*_*y*_). Interpenetrating phases satisfy

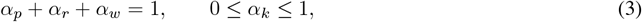

and, under phasewise incompressibility,

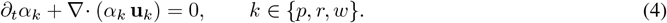

Phase momenta evolve as

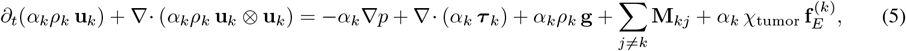

with a common mixture pressure *p*, viscous stress ***τ*** _*k*_, and interphase exchange **M**_*kj*_ satisfying Σ_*k*_Σ_*j* ≠ *k*_ **M**_*kj*_ = **0**. Phase densities used here are *ρ*_*p*_ = 1030 kg*/*m^3^, *ρ*_*r*_ = 1100 kg*/*m^3^, *ρ*_*w*_ = 1080 kg*/*m^3^ [Nithiarasu, 2022, Schwartz, R. S. and Conley, C. L., 2020]. The indicator *χ*_tumor_ ∈ { 0, 1} activates electrohydrodynamic (EHD) forcing only in the tumor–vessel subdomain.

The electroquasistatic potential satisfies

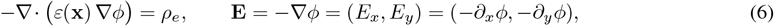

where *ε* = *ϵ*_0_*ϵ*_*r*_ is the permittivity (*F/m*), *ϵ*_0_ is the vacuum permittivity, *ϵ*_*r*_ is the case-specific relative permittivity, and *ρ*_*e*_ is the free charge density (*C/m*^3^). The Kelvin (electrohydrodynamic) body force (**f**_*E*_) added to Eq.5 is

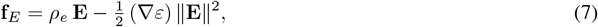

as in continuum EHD theory [Landau et al., 2013, Saville, 1997, Castellanos et al., 2003]. In this study *ε* is treated as spatially uniform unless otherwise noted, so **f**_*E*_ = *ρ*_*e*_ **E**. The Fluent UDF applies the phase-aggregated force components (where *E* is the electric potential)

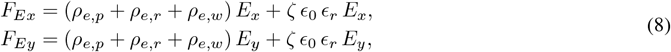

where *ρ*_*e,k*_ are phase charge densities and *ρ*_*e*_ = ∑_*k*_ *ρ*_*e,k*_; *ζ* is the zeta potential applied on EHD-active glycocalyx patches. Electrical properties used in Eq.6–8 are: *ζ* = − 0.01 V [Squires and Bazant, 2006]; *ϵ*_0_ = 8.854 × 10^−12^ F*/*m; phase conductivities *σ*_*p*_ = 1.2 S*/*m [Pauly and Schwan, 1966], *σ*_*r*_ = 0.52 S*/*m [Zhbanov and Yang, 2015], *σ*_*w*_ = 0.64 S*/*m [Yang et al., 1999]; and a case-specific *ϵ*_*r*_ stated with each EHD case.

The blood’s shear-thinning behavior is represented by [Jung and Hassanein, 2008, Jung, J. et al., 2006]. With hematocrit *α*_*r*_ (RBC volume fraction) and scalar shear rate 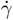,

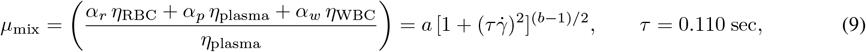

where *µ*_mix_ is the mixture dynamic viscosity (Pa·s), *η*_plasma_ = 0.001 kg*/*(m · s) [Nithiarasu, 2022], and we take *η*_WBC_ = 0.011 kg*/*(m · s) [Schwartz, R. S. and Conley, C. L., 2020]. The effective RBC contribution is captured through the hematocrit-dependent mixture law; no separate constant *η*_RBC_ is prescribed. The coefficients *a*(*α*_*r*_) and *b*(*α*_*r*_) are dimensionless Carreau–Yasuda–type fits to *α*_*r*_:

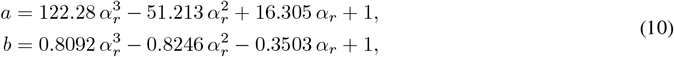

and, for 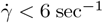,

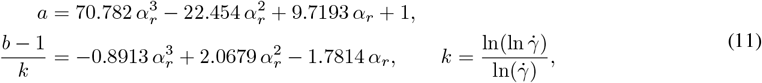

where *k* is the (dimensionless) Yasuda exponent. Interphase drag follows Schiller–Naumann [Naumann and Schiller, 1935]; RBC agglomeration enters via the dynamic shape factor

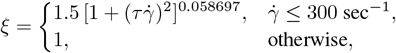

while WBCs use *ξ* = 1. Characteristic cell sizes used in closures are *d*_RBC_ = 7 *µ*m and *d*_WBC_ = 14 *µ*m [Nithiarasu, 2022, Schwartz, R. S. and Conley, C. L., 2020].

Boundary conditions for the tumor-vessel calculation are: pressure inlet *p*_in_ = 3325 Pa and pressure outlet *p*_out_ = 2128 Pa [Zhao, G. et al., 2007]. At the outlet, the backflow volume fraction is set to RBC= 1 and WBC= 0 to prevent unphysical WBC re-entry while remaining consistent with the dispersed-phase specification. Electric potentials *ϕ* are applied only on tumor–vessel boundaries where EHD is active; elsewhere *∂*_*n*_*ϕ* = 0. Time integration employed a constant time step Δ*t* = 1.0 × 10^−4^ sec; each vessel case was advanced for 2000 iterations, matching the ECM simulation, with convergence monitored via mass and momentum residuals and outlet flux balances. Computations were run in parallel on four Xeon cores at 3.1 GHz. From each of the fifteen vessel geometries, plasma pressure and velocity were sampled at the fenestra-center location (wall depression); the arithmetic averages of these fields define the inlet data subsequently used in the ECM (2D) plasma-perfusion simulation.

#### 2.2.2 Simulation method for plasma perfusion inside the ECM tumor region

Plasma perfusion in the ECM domain was modeled as viscous–laminar, transient, two dimensional flow using the Eulerian multiphase formulation in ANSYS FLUENT 2023 R1 [Afolabi, E. A. and Lee, J., 2014]. A shared pressure field is solved while phasewise continuity and momentum equations advance the primary (plasma) and secondary (air) phases. Interphase coupling is represented through drag exchange terms with coefficients evaluated from the local Reynolds number, consistent with the Eulerian framework [Afolabi, E. A. and Lee, J., 2014]. Electrohydrodynamic forcing is not applied in the ECM calculation (EHD is restricted to the vessel domain), so the momentum balances here contain only hydrodynamic and interphase exchange terms. The ECM inlet condition is constructed from the tumor vessel simulations at a fixed snapshot in time. For each of the *N* = 15 vessel cases, plasma pressure *p*^(*i*)^(*x, y, t*) and velocity **u**^(*i*)^(*x, y, t*) were sampled at the fenestra-center location at *t* = Δ*t* = 1.0 × 10^−4^ sec. The cross-model averages

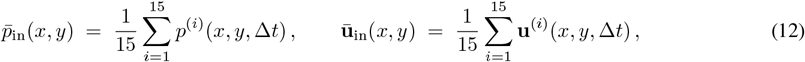

define the prescribed inlet pressure and velocity for the ECM inflow boundary, ensuring consistency with the three-phase vessel-scale plasma transport trend. To create a clean plasma-advance front in the interstitial spaces, the secondary phase was taken as air. The ECM domain was initialized as air-filled (air volume fraction *α*_air_ = 1, plasma volume fraction *α*_plasma_ = 0), and plasma displaced air over time under the inlet fields Eq.12. The tumor slice geometry has three outlets; each outlet was set to a fixed pressure of 2780 Pa, following prior studies [Wu, M. et al., 2013] and consistent with elevated interstitial pressures reported in tumors [Sven, K. and Josipa, F., 2007, Stylianopoulos et al., 2018b, Heldin, C. et al., 2004]. No-slip conditions were enforced on all solid boundaries (fiber and ECM walls). ime integration used a constant time step Δ*t* = 1.0 × 10^−4^ sec; each ECM run was advanced for 2000 iterations (matching the tumor vessel simulation), with convergence assessed by residual reduction for mass and momentum and by outlet flux balances. Computations were executed in parallel on four Xeon cores at 3.1 GHz with a typical wall-clock time of 3–4 h. Material properties matched those used elsewhere in this study: plasma density and viscosity *ρ*_p_ = 1030 kg*/*m^3^ and *η*_p_ = 0.001 kg*/*(m · s) [Nithiarasu, 2022]; air density and viscosity *ρ*_air_ = 1.225 kg*/*m^3^ and *η*_air_ = 1.8 × 10^−5^ kg*/*(m · s) [Porterfield, W. W. and Kruse, W., 1995]. These settings, together with the inlet fields 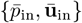 and outlet pressures, define a closed initial–boundary-value problem for plasma permeation through the ECM microstructure, from which we compute the space–time evolution of plasma volume fraction, interstitial velocities, and cumulative perfusion metrics.

### 2.3 Experimental setup

To replicate the physical features of a tumorous microenvironment (same CAD model of Fig.2E), we fabricated a biomimetic pillar array using a Form 3+ stereolithography (SLA) resin printer (Formlabs Inc., USA). The structure (Fig.3C) was printed in clear photopolymer resin, achieving a spatial resolution of 50*µ* m. The design matched the packing fraction of 0.30 observed within tumors. After printing, the structure was rinsed in isopropyl alcohol and post-cured under UV light. The flow through the pillar network was driven using red-dyed deionized water as the working fluid. The fluid was supplied at a constant volumetric flow rate using a Harvard Apparatus syringe pump (Fig.3B, enabling precise control of perfusion conditions within the range of 1 − 3 *mL/min* (Fig.3D-E *are for 2 mL/min*). This range ensured laminar flow within the micro-pillared structure and preserved quasi-two dimensional dynamics consistent with the assumptions of the numerical models. The transparent resin structure was illuminated from below using a diffuse white light source, while the evolving liquid front was imaged from above with a digital camera. The recorded image sequences were post-processed using image analysis software to extract the time evolution of the wetted area. Binarized frames were used to calculate the area of liquid penetration as a function of time, from which the spreading dynamics were quantified.

**Figure 3:**
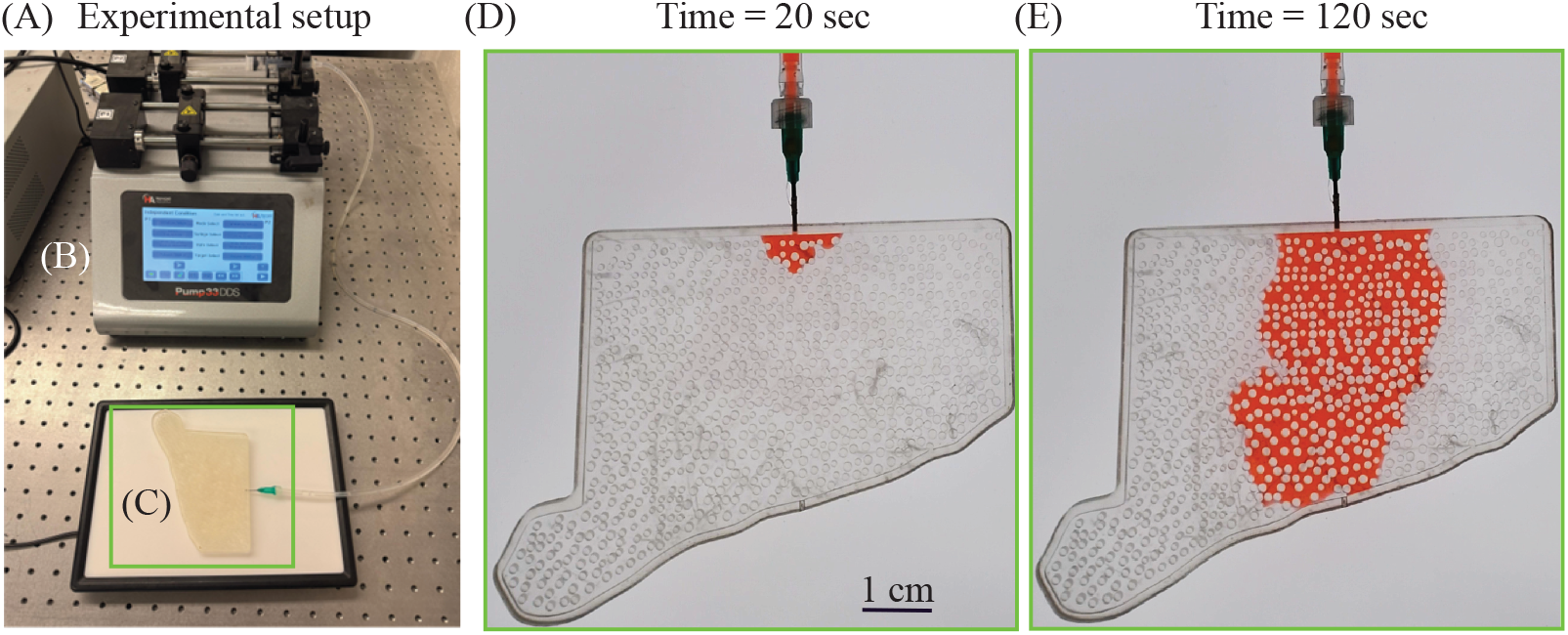
Microfluidic visualization of dye-driven perfusion through a 3D-printed tumor ECM: (A) Experimental setup for the microfluidic test. Red–dyed deionized water was used to mimic plasma perfusion through a tumor ECM domain 3D-printed in clear photopolymer resin (C, green box). (B) Harvard Apparatus syringe pump delivering a constant 2 *mL/min* flow. (D–E) Time-lapse percolation within the ECM at 20 sec and 120 sec, respectively, showing propagation of the dyed front inside the ECM; the green frame indicates the zoomed field of view from (C) during flow. Scale bar: 1 cm.

### 2.4 CFD-informed reverse advection-diffusion (RAD) model

Plasma-borne solute transport in solid tumors reflects a shifting balance between advection and diffusion, but elevated interstitial fluid pressure and dense ECM packing can suppress forward flow and induce reverse advection—migration opposite to the bulk direction. Because Darcy’s law and the Starling formulation do not capture this local reversal or geometric obstruction [Baish et al., 2011, Sugihara-Seki and Fu, 2005], we introduce a reverse advection–diffusion (RAD) model that couples theoretical transport to a CFD-informed advection field at the fenestra derived from multiphase simulations.

First, a multiphase numerical simulation (Sec. 2.2.1) is performed to characterize the vessel-to-ECM entry and to extract the inlet advection profile, i.e., the *y*-directed average velocity at the fenestra used as *ω*(*y*) in Eq.13. The RAD model is posed on a reduced-order rectangular ECM domain of size 10 × 5 *µ*m^2^ with a 0.3 *µ*m fenestra through which plasma enters. A total of 40 circular fiber inclusions are placed with minimum spacing (no overlap) to maintain a packing fraction *ϕ* = 0.30 [Andersson et al., 2008, Nier et al., 2016, Spatarelu et al., 2019, Yeasin, 2025], yielding an average fiber radius 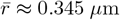. The geometry is reconstructed in Wolfram Mathematica and meshed via ToElementMesh into an unstructured triangular grid (~ 5.5 × 10^4^ elements), providing sufficient resolution near the fenestra and fiber boundaries.

The governing transport equation is

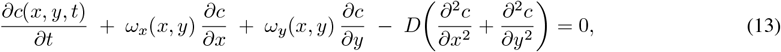

where *c*(*x, y, t*) is solute concentration, *D* is the effective diffusivity, and ***ω***(*x, y*) = (*ω*_*x*_, *ω*_*y*_) is the imposed advection field; in practice we take *ω*_*x*_ = 0 and prescribe *ω*_*y*_ from the CFD-derived inlet profile (uniform along the fenestra in Sec.2.2.1). We initialize with *c*(*x, y*, 0) = 0 and apply a constant inlet concentration of 8.0 × 10^13^—twelve orders of magnitude above the physiological protein concentration in plasma (~ 80 mg*/*mL [Leeman et al., 2018])—at the fenestra. To match CFD timescales in this reduced domain, the inlet concentration is scaled (constant factor), while the effective diffusivity is set to *D* = 1270 *µ*m^2^*/*s [Muraoka et al., 2008]. Zero-flux (Neumann) boundary conditions are enforced on lateral and outlet boundaries to allow convective outflow without artificial back-diffusion, and all fiber surfaces are treated as impermeable (**n**·∇*c* = 0).

For validation on the same simplified geometry, we conducted a two phase numerical simulation in ANSYS Fluent 2023 R1. The domain was constructed in ANSYS DesignModeler 2023 R1 and discretized with an unstructured triangular mesh (approximately 7.1 × 10^4^ elements) with local refinement near the fenestra and along the patched-wall segments. To accommodate the concentration inlet prescribed by the theoretical model, only the outlet boundary condition was modified for this validation case; all other settings (solver options, material properties, numerics) were kept identical to Sec. 2.2.2. Geometry and boundary placements were identical to those used for the RAD domain.

## 3 Results

### 3.1 Comparison of plasma perfusion from vessel to ECM with and without EHD

We quantify how electrohydrodynamics (EHD) in the tumor vessel modifies the plasma delivered through the fenestra into the ECM, relative to an otherwise identical non-EHD baseline. Using the representative geometry (Model 5), we select the same region of interest (ROI)–the fenestra –in both simulations (Fig. 5A–B). Pixel colors in the ROI are mapped to a dimensionless plasma intensity *S* ∈ [0, 1] producing two paired empirical distributions over the identical ROI: *S*_EHD_ and *S*_non_-_EHD_. The histogram comparison in Fig. 5E shows a decisive rightward shift with EHD. Denote the sample means by 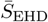 and 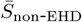, and standard deviations by *s*_EHD_ and *s*_non_-_EHD_. and let *s*_*p*_ be the pooled standard deviation. The standardized mean difference (Cohen’s *d*) [Cohen, 2013] is

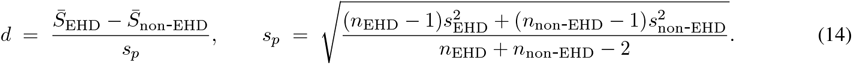

**Figure 4:**
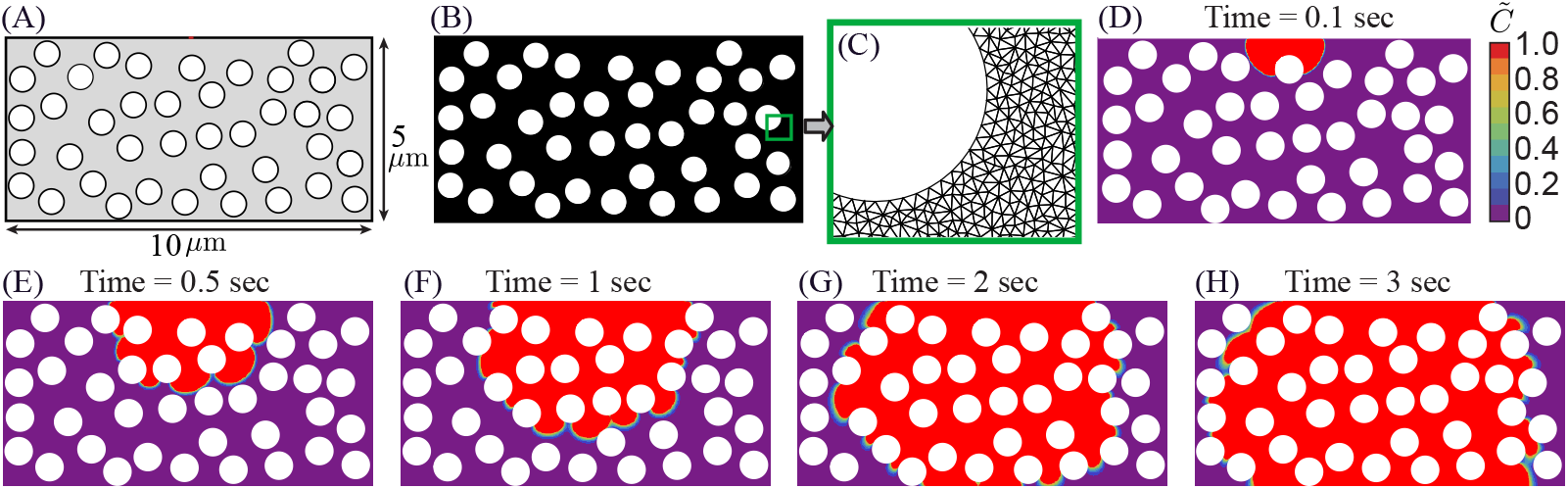
CFD-informed reverse advection-diffusion RAD model showing solute plume evolution through a reduced-order fiber-dense tumor ECM: (A) The simplified rectangular computational domain (10 × 5 *µ*m^2^) used for the reduced-order RAD model, containing randomly distributed circular ECM fiber bundles occupying 30% of the rectangular domain area. All fibers have equal radius and act as impermeable structures, mimicking dense tumor stroma and restricting solute motion. (B) The high-resolution unstructured triangular finite-element mesh generated for the theoretical analysis. Whereas (C) provides a zoom view of the mesh in the fenestra region, highlighting refined discretization around fiber boundaries to resolve steep concentration gradients. (D-H) The predicted spatiotemporal evolution of plasma-borne solute under the advection-diffusion transport model at *t* = 0.1 sec, 0.5 sec, 1.0 sec, 2.0 sec, and 3.0 sec, respectively, where 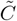 represent the normalized concentration field.

**Figure 5:**
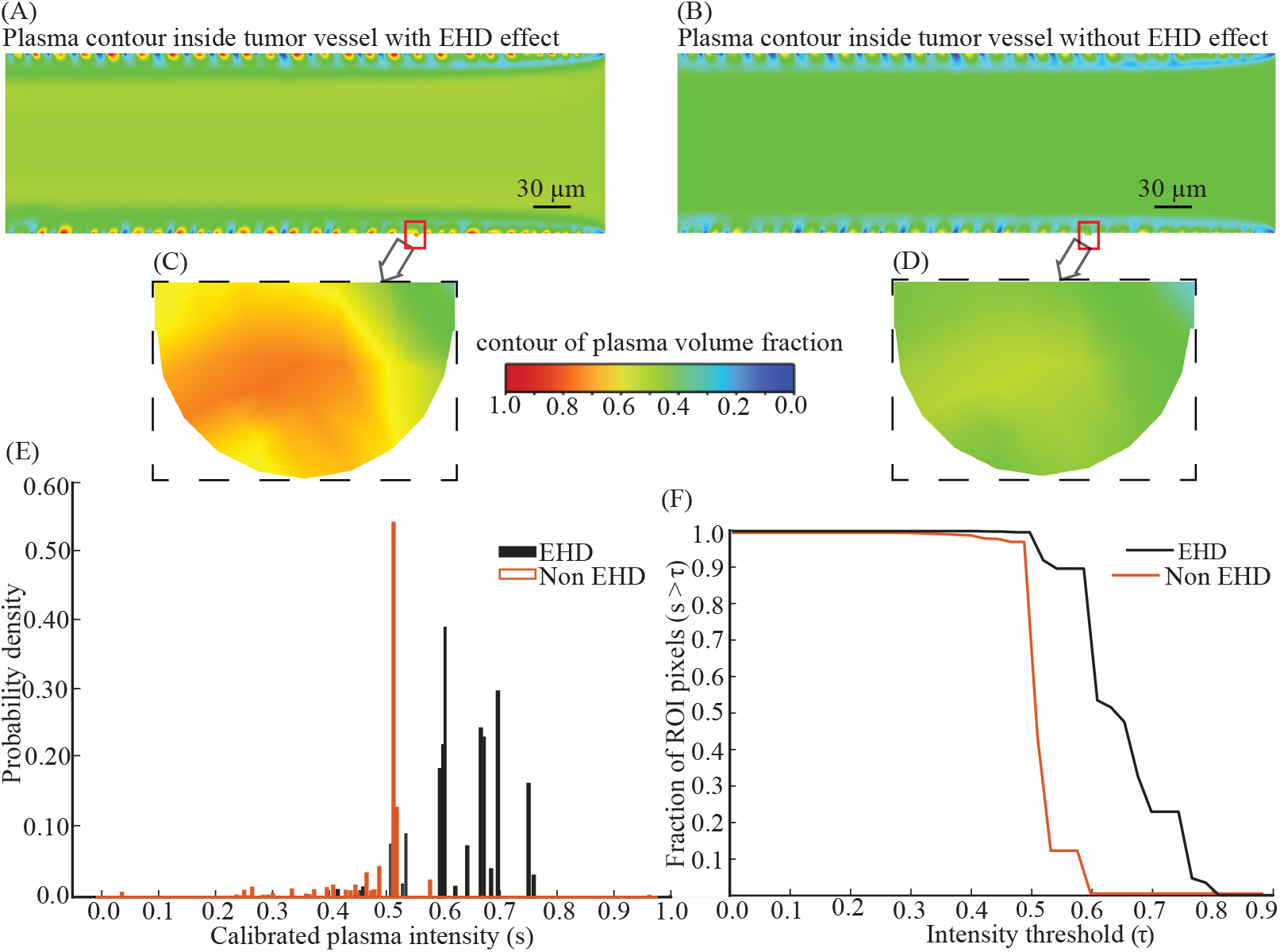
EHD flow yields enhanced, more persistent plasma penetration at the fenestra compared with non-EHD flow: (A-B) Plasma (primary phase) contour inside the tumor vessel with EHD (A) and without EHD (B). (C–D) Zoomed fenestra ROI from panels A and B (same simulation: Model 5). The dashed rectangle delineates the identical physical area used for pixel counting and threshold analysis. (E) Fenestra-ROI intensity distributions show a clear rightward shift under EHD with minimal overlap, indicating stronger local plasma penetration; mean intensities: 0.650 (EHD) and 0.525 (non-EHD). (F) Threshold (survival) curves summarize robustness across cutoffs: the EHD curve dominates across thresholds, and the area-between-curves (ABC) equals 0.125, demonstrating more persistent high-intensity pixels only with EHD.

For Model 5, 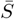 increases from 0.525 (non–EHD) to 0.650 (EHD), yielding *d* = 2.05 (“very large” separation). A threshold-free dominance metric is the probability of superiority (AUC),

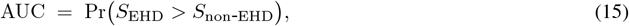

from which Cliff’s delta follows as

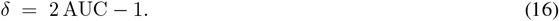

Here, AUC = 0.902 and *δ* = 0.804 indicate minimal overlap between conditions. A robust location shift is summarized by the Hodges–Lehmann median difference,

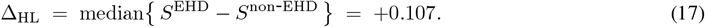

To assess areal coverage at clinically relevant intensities, we examine the exceedance (survival) curves (Fig.5F) over a threshold *τ* ∈ [0, 1]:

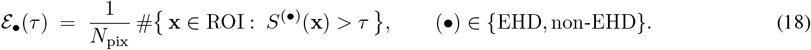

Fig. 5F shows *ℰ*_EHD_(*τ*) (black) lies strictly above *ℰ*_non_(*τ*) (orange) across the inspected range, i.e., larger high-intensity area for any chosen cutoff. The area between curves (ABC) provides a cutoff-agnostic summary:

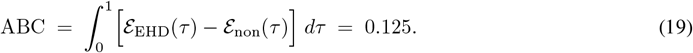

At a stringent threshold of *τ* = 0.7, the high-intensity fraction is 22.92% (EHD) versus 0.0% (non-EHD), clearly showing a strongly perfused area under EHD. In Fig.5, panels E–F jointly indicate that EHD (via patched, glycocalyx-mediated wall forcing) increases both how much of the fenestra area attains high plasma fraction (larger exceedance across *τ*) and how strongly it is filled (right-shifted intensity distribution). These inlet-side gains are exactly the mechanism that yields the faster and broader percolation observed downstream in the ECM domain (Sec. 3.2). Across all 15 models, Fig. 6A–B demonstrates that the mean calibrated intensity under EHD is consistently higher than under Non–EHD. Fig6A shows this pattern model–by–model, while panel B summarizes the across–model effect: grand means of 0.722 (EHD) versus 0.576 (Non–EHD), with a mean improvement Δ = 0.146 (95% CI [0.124, 0.166]; Wilcoxon signed–rank *p* = 6.1 × 10^−5^). Our objective here is to establish the condition effect (EHD vs. Non–EHD), not to optimize fenestra placement; accordingly, we do not rank or compare the EHD models against one another by location.

**Figure 6:**
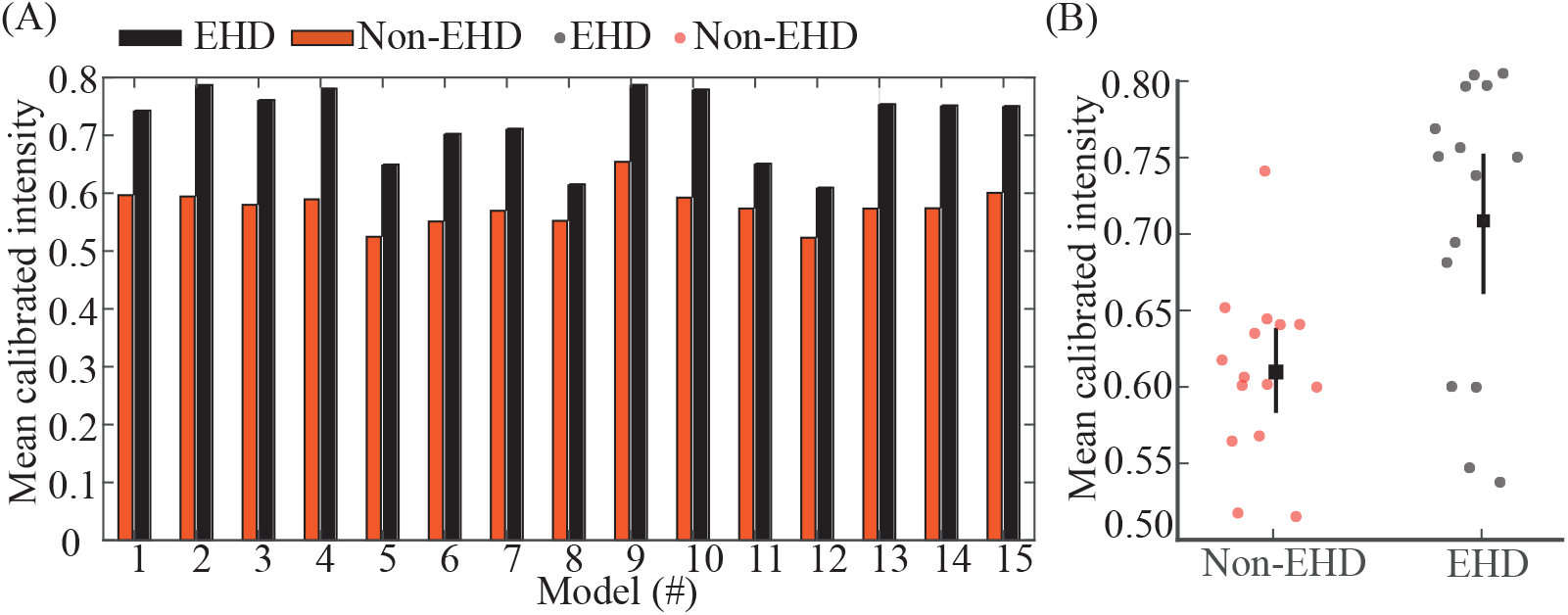
EHD flow consistently yields higher mean plasma penetration than non-EHD across all 15 models: (A) For Models 1–15 (fenestra center translated from near inlet to near outlet), grouped bars compare Non–EHD and EHD mean calibrated intensity for each model; higher values indicate greater plasma penetration. EHD exceeds Non–EHD in every model. (B) Per–model means (dots) with grand mean and 95% bootstrap CI (square with whiskers) for each condition. Across models, mean Non–EHD = 0.576 and mean EHD = 0.722 (mean Δ = 0.146, 95% CI [0.124, 0.166]; Wilcoxon signed–rank *p* = 6.1 × 10^−5^).

### 3.2 Plasma percolation inside the tumor ECM domain

With the tumor-domain inlet fields 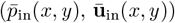 constructed from the *N* = 15 vessel simulations (Sec. 2.2.2, Eq.12), the flow advances the air–plasma interface through the tumor ECM domain. The spatiotemporal evolution exhibits a coherent percolation pattern: an early thin, inlet-aligned jet penetrates along preferential low-resistance pathways; intermediate times show lateral broadening and progressive saturation of interstitial voids; and late times approach a nearly uniform, high–plasma-fraction interior behind the front. This progression is visualized in Fig. 7A–D via contours of local plasma volume fraction, where red denotes the advancing high-fraction front, yellow/green the graded rim, and blue the not-yet-perfused interstitium. To quantify front advance and rim thickening, we compute an image-based color occupancy ratio for each frame by normalizing the blue (unfilled) pixels to *B* = 1 and reporting the triplet *R* : (*G*+*Y*) : *B* after excluding white pores from the counts. A rise in *R* : *B* signals deeper filling, and a rise in (*G*+*Y*) : *B* signals a thickening front. As summarized in Table 2, *R* : *B* grows monotonically from ≈ 2 × 10^−3^ at *t* = 0.01 sec to ≈ 6.83 at *t* = 0.20 sec, with a concurrent increase in (*G*+*Y*) : *B*. Consistent with the visual maps, early times are advection-dominant (small *R* : *B*, thin inlet-aligned plume); by mid-times the rim thickens as cross-stream spreading increases; at late time most accessible voids are filled (blue nearly exhausted) and the field behind the front approaches homogeneous red.

**Table 2:**
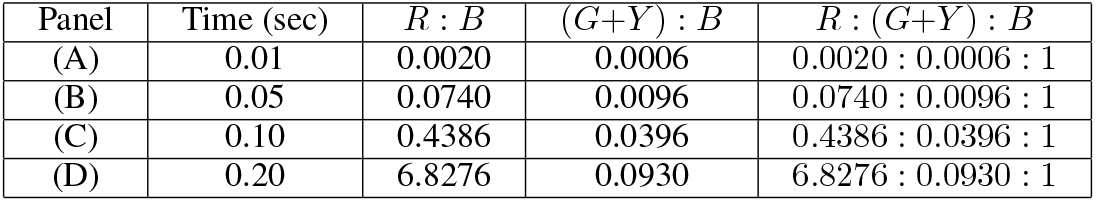
Color-occupancy ratios for Fig.7 (A–D). Ratios are computed from pixel counts with blue normalized to 1; white pore pixels are excluded. Increasing *R* : *B* and (*G*+*Y*) : *B* indicate deeper penetration and a thickening front.

| Panel | Time (sec) | $R : B$ | $(G+Y) : B$ | $R : (G+Y) : B$ |
| --- | --- | --- | --- | --- |
| (A) | 0.01 | 0.0020 | 0.0006 | 0.0020 : 0.0006 : 1 |
| (B) | 0.05 | 0.0740 | 0.0096 | 0.0740 : 0.0096 : 1 |
| (C) | 0.10 | 0.4386 | 0.0396 | 0.4386 : 0.0396 : 1 |
| (D) | 0.20 | 6.8276 | 0.0930 | 6.8276 : 0.0930 : 1 |

**Figure 7:**
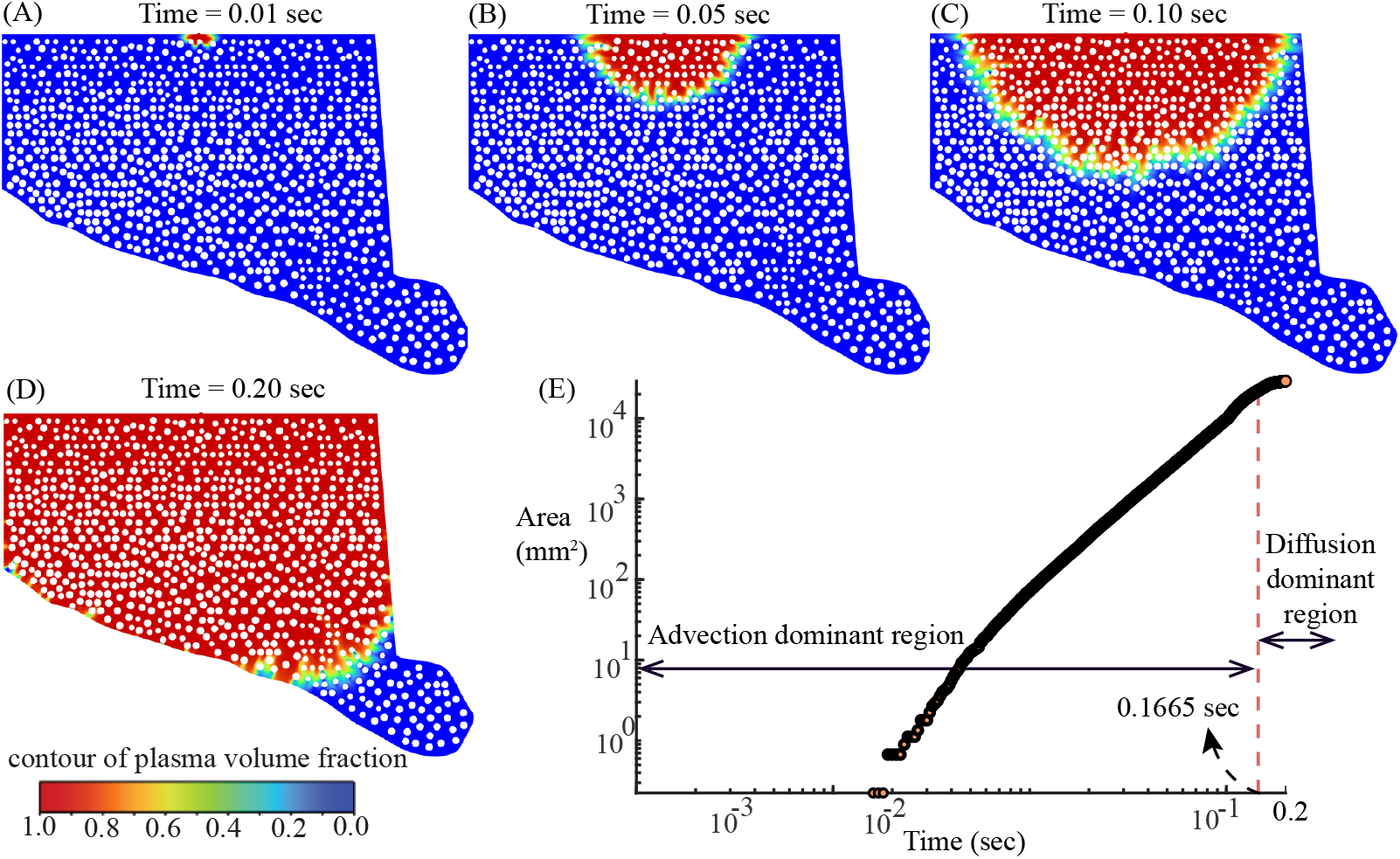
Spatiotemporal spreading of plasma in the tumor ECM shows an advection-to-diffusion transition in wetted area growth: (A–D) Contours of plasma volume fraction at (A) 0.01 sec, (B) 0.05 sec, (C) 0.10 sec, and (D) 0.20 sec. White circles denote impermeable fibers; blue is unfilled interstitium. The 0–1 color bar maps local plasma fraction. The wetted region expands from a thin inlet plume to a broad front, illustrating a qualitative shift from advection-driven advance toward diffusion-controlled spreading. (E) Wetted area versus time on log–log axes. The red dashed line marks the transition time *t*\* = 1.665 × 10^−1^ sec, separating the advection-dominant region (left) from the diffusion dominant region (right).

A complementary, domain–integrated metric is the wetted area *A*(*t*) of non–blue pixels. On log–log axes we analyze (*x*_*i*_, *y*_*i*_) = (log_10_ *t*_*i*_, log_10_ *A*_*i*_) so that a power law *A* ~ *t*^*α*^ appears as a straight line:

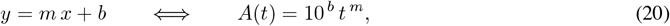

with exponent *α* = *m*. A two–segment least–squares fit in (*x, y*) identifies a break point *t*\* that minimizes the total residual sum of squares. Using this procedure we obtain *t*\* = 0.1665 sec, which partitions the record into advection region: t ∈ [1.60 × 10^−3^, 1.664 × 10^−1^] sec, and diffusion dominant region: t ∈ [1.666 × 10^−1^, 2.00 × 10^−1^] sec. Within any 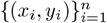, ordinary least squares gives (see the Table3)

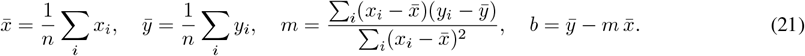

**Table 3:** Segment-wise fits on (log_10_ *t*, log_10_ *A*) for log_10_ *A* = *m* log_10_ *t* + *b*.

| Segment | $t$-range (sec) | $m$ | $b$ | $R^2$ |
| --- | --- | --- | --- | --- |
| Advection region | $[1.60 \times 10^{-3}, 1.664 \times 10^{-1}]$ | 2.21 | 5.23 | 0.99 |
| Diffusion dominant region | $[1.666 \times 10^{-1}, 2.00 \times 10^{-1}]$ | 0.41 | 3.75 | 0.98 |

To summarize the transition at *t*\*, we report the one–sided model predictions,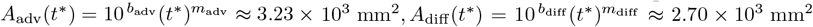. The advection side extrapolation is therefore higher at the breakpoint (offset ≈ 20% relative to the diffusion-side value; ≈ 18% relative to the mean of the two), which is typical for an unconstrained two–line fit and indicates an abrupt change in kinetics rather than a perfectly continuous crossover. The effective exponent drops from super–linear to subunit (*m* : 2.21 to 0.41), consistent with a shift from advection–driven growth to a diffusion–controlled late regime; this interpretation agrees with the RGB composition (persistent graded rim).

### 3.3 Experiment–CFD validation of wetted-area growth

Top view images (Fig.3D-E) from the experiments were segmented to obtain wetted pixel counts over time and converted to physical area, yielding *A*(*t*) in Fig. 8E. The experiment exhibits advection dominated growth overall, with an early shoulder consistent with *A*(*t*) ~ *t*^1*/*2^ before a clear linear trend *A*(*t*) ~ *t* becomes established; we identify the knee at 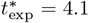 sec. For a like-for-like benchmark, we ran a separate 2D CFD configuration (distinct from Sec. 2.2.2) using a pressure-driven inlet to match the experimental 2 mL*/*min; plasma and air properties were unchanged, and the front evolution appears in Fig. 8A–C. Applying the same area-extraction to the simulation yields the same qualitative behavior; advection-dominated growth with a brief early shoulder—and a much earlier knee at 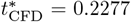 sec (Fig. 8D). Thus, both experiment and CFD are advection-dominated; the primary discrepancy is the transition timescale (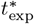 versus 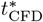), which we analyze in Sec. 4.2 in terms of inlet start-up/wetting effects and geometric idealization.

**Figure 8:**
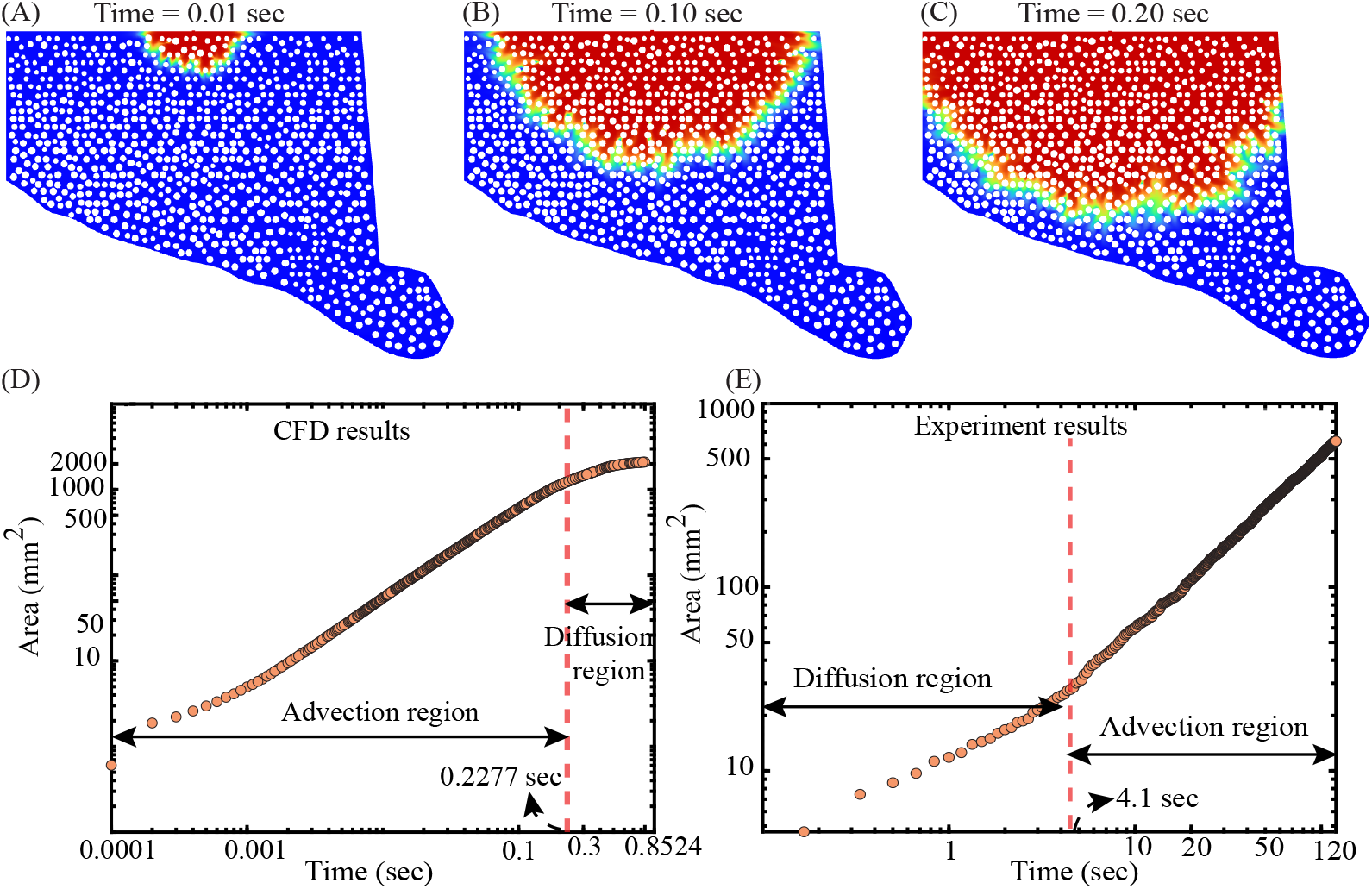
CFD and experiments both show advection-dominated growth regimes in ECM wetting, with different transition times: (A–C) Snapshots show front advance at *t* = 0.01, 0.10, 0.20, sec respectively. (D) log–log growth of area *A*(*t*) with an advection-dominated regime transitioning to diffusion at 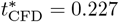 sec. (E) early diffusion-dominated growth (*A* ~ *t*^1*/*2^) transitioning to advection (*A* ~ *t*) at 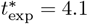 sec.

### 3.4 Comparison of theoretical model predictions with corresponding numerical simulation outcomes

The reverse advection–diffusion (RAD) model reproduces the same two-stage transport seen in the like-for-like CFD of the simplified rectangular domain: an inlet-aligned, advection-driven entry plume followed by diffusion-dominated homogenization as the interior fills (theory snapshots in Fig. 4D–H; CFD snapshots in Fig. 9A–D). To quantify kinetics, we track the occupied interstitial area *A*(*t*) for both models on log–log axes (Fig. 9E–F). In each case the slope transitions from a steep, advection-driven regime to a shallow, diffusion-controlled regime, with breakpoints at 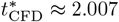 sec (Fig. 9E) and 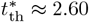 sec (Fig. 9F).

**Figure 9:**
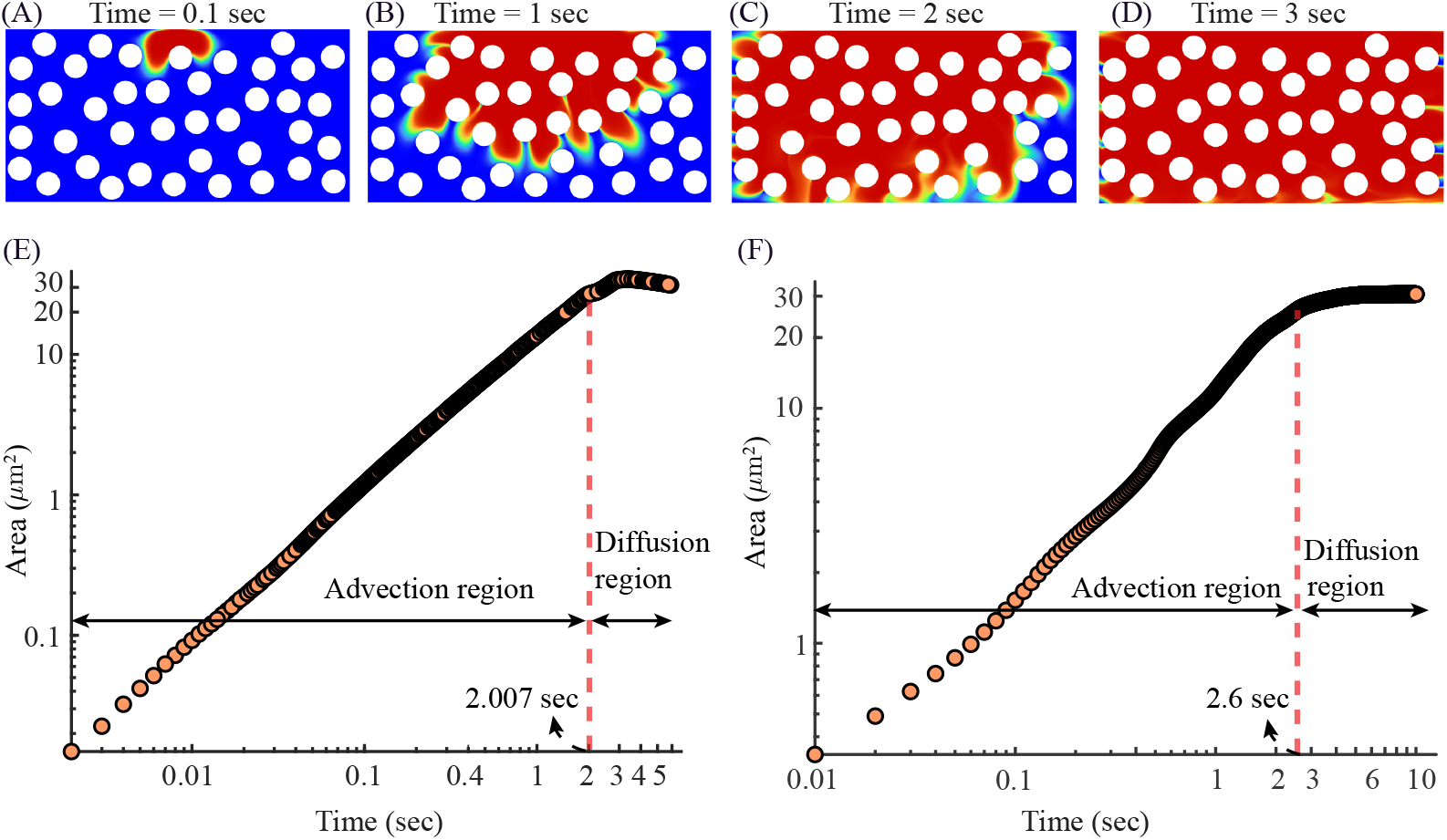
Comparison of CFD and RAD model for plasma transport transition: (A) shows the CFD-predicted plasma propagation at *t* = 0.1, 1, 2, and 3 sec, respectively. Plasma first penetrates swiftly along preferential pathways between fibers, then progressively spreads laterally and fills the matrix, reflecting a transition from advection-dominated entry to diffusion-driven equilibration. (E) shows the occupied interstitial area *A*(*t*) as a function of time for the CFD solution, demonstrating a steep early-time growth regime followed by a slowed diffusion-controlled regime as the domain fills. (F) shows the corresponding theoretical RAD model prediction for *A*(*t*), showing the same trend as the CFD solution.

**Figure 10:**
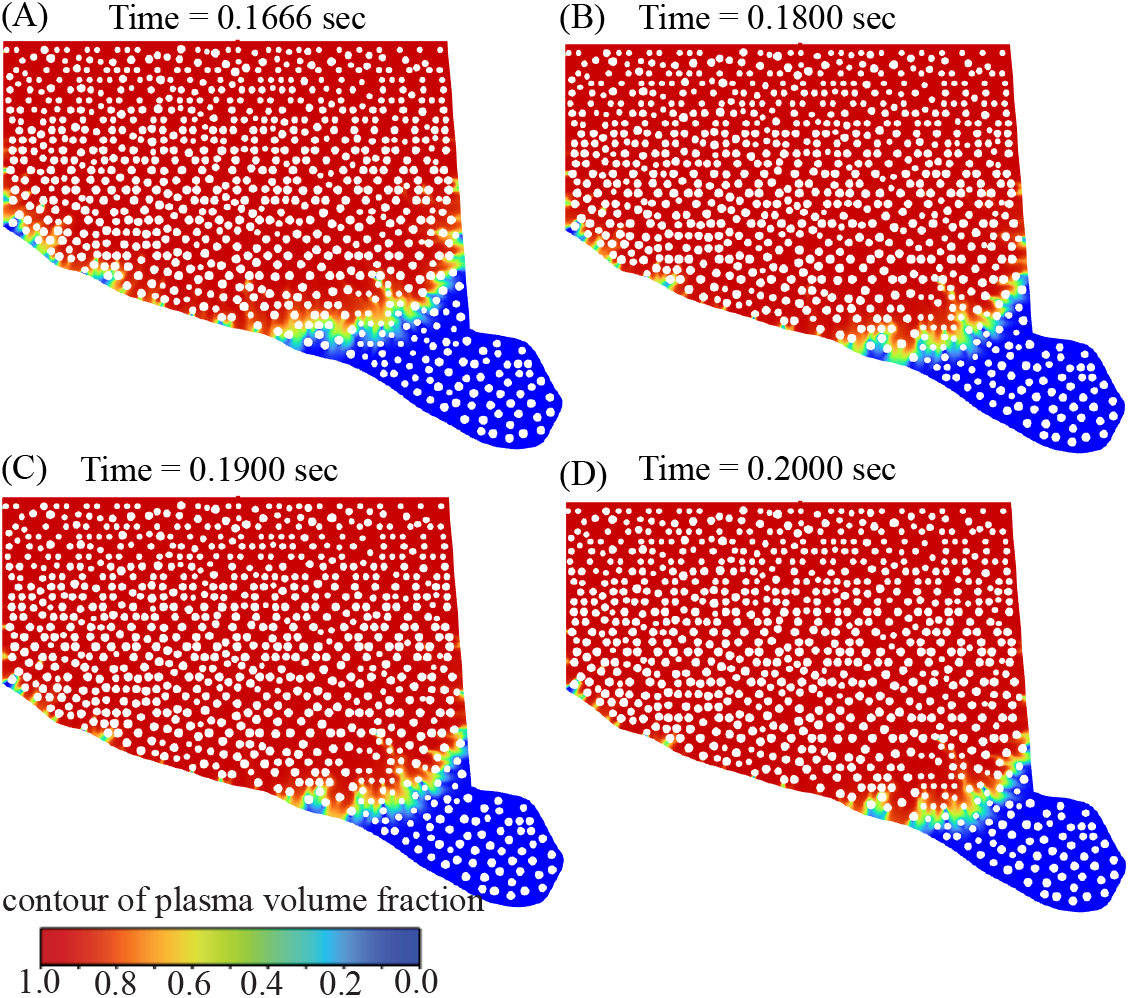
Temporal evolution of plasma penetration inside tumor ECM domain at different flow times: Panels show snapshots of plasma perfusion inside the tumor domain at (A) t=0.1666 sec, (B) 0.1800 sec, (C) 0.1900 sec, and (D) 0.2000 sec within the diffusion-dominated region. Red denotes the high-fraction bulk, green+yellow denote the graded rim, and blue denotes the unperfused region. Panels show a persistent graded rim leading a slowly advancing bulk with a gradual recession of blue—indicates gradient-limited (diffusive) penetration through a tortuous, fiber dense matrix, consistent with subunit area-growth scaling.

We summarize the post-transition timescale offset by the simple ratio

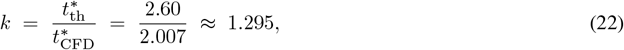

indicating that the theoretical curve evolves about 1.295 times more slowly than the CFD result on this domain. If we rescale the theoretical time axis by this factor,

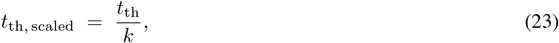

it will bring the *A*(*t*) trajectories into close agreement in both the early advection regime and the late diffusion-dominated regime. Thus, the reduced-order RAD model captures the same mechanistic shift observed in CFD and, after a single temporal calibration, serves as an accurate and computationally efficient surrogate for fully resolved flow simulations on this geometry.

## 4 Discussion

### 4.1 Diffusion dominant late regime with subunit area scaling

In the late regime (*t > t*\* = 0.1665 sec)), wetted–area growth is governed by diffusion even though the log–log exponent is subunit (*m*_diff_ ≈ 0.41). A persistent graded rim (green+yellow) leading the bulk (red), quantified by slowly varying color ratios (Table 4), is consistent with diffusion–limited penetration through a tortuous, fiber–dense matrix. In diffusion–limited flow, the characteristic penetration length obeys 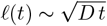, so observables that scale with length (e.g., a band–like front across a roughly fixed width) grow as *t*^1*/*2^, not linearly [Crank, 1979]. Consequently, the area can grow sublinearly (*m <* 1) and still be diffusion–controlled. In ideal band–like geometries one expects *A*∝ *t*^1*/*2^ (*m* = 0.5); values below 0.5 are expected in complex matrices where tortuosity, dead–ends, or partial channelization slow the effective front (subdiffusion) [Sahimi, 2011, Metzler and Klafter, 2000, Berkowitz et al., 2000, Edery et al., 2014].

**Table 4:**
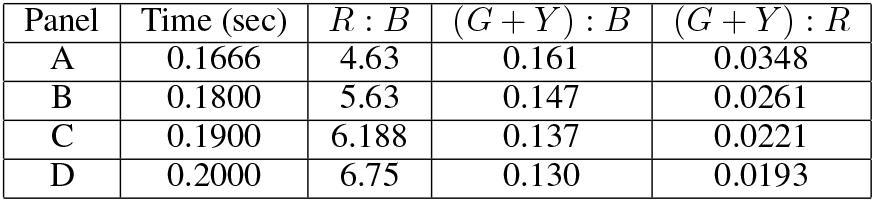
Color–composition ratios in the diffusion–dominant region in Fig. 10.

Independent image–based composition metrics corroborate a diffusion–controlled tail. Post *t*\* snapshots show a persistent graded rim (green+yellow) leading the high–fraction bulk (red) while the unperfused region (blue) depletes gradually. Quantitatively, the *R* : *B* increases only slowly from 4.63 to 6.75 between t=0.1666 and 0.2000 sec, and the (*G*+*Y*) : *B* remains appreciable (0.161 to 0.130). Crucially, the rim–to–bulk ratio ((*G*+*Y*) : *R*) stays non-vanishing and declines only mildly ~ 3.5% → 1.9%, indicating a broad, gradient–limited front rather than a sharp advective discontinuity at high Péclet number.

### 4.2 On the experiment–CFD timescale discrepancy and thickness scaling

We first quantify the temporal discrepancy by comparing the post-transition analysis. The corresponding timescale ratio is

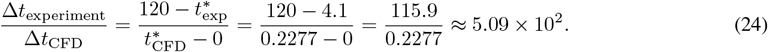

This result indicates that the experimental timescale is approximately 509 times longer than the computational timescale, emphasizing the profound effect of physical constraints present in the 3D experimental setup but absent in the 2D numerical model.

Under the constant–flow (pump–driven) conditions of the experiment, mass conservation in a thin layer of thickness *H* gives 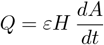, so the characteristic time obeys Δ*t* ∝ *H* at leading order. Interpreting the 2D CFD as a per–depth surrogate of effective thickness *h*_eff_, the transition–time ratio therefore scales as

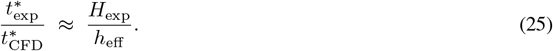

With *H*_experiment_ = 7.5 mm = 7500 *µm* and the measured ratio ≈ 1.85 × 10^2^, we infer

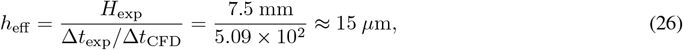

which explains the *𝒪* (10^2^) gap directly from the 3D–to–2D thickness difference. To reconcile the curves graphically (Fig. 11), we rescale CFD time and area by

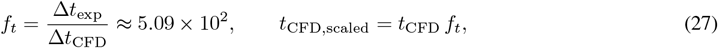

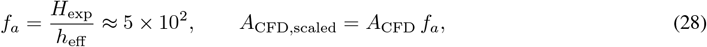

and plot the residual as

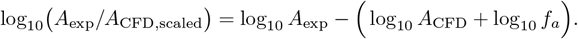

**Figure 11:**
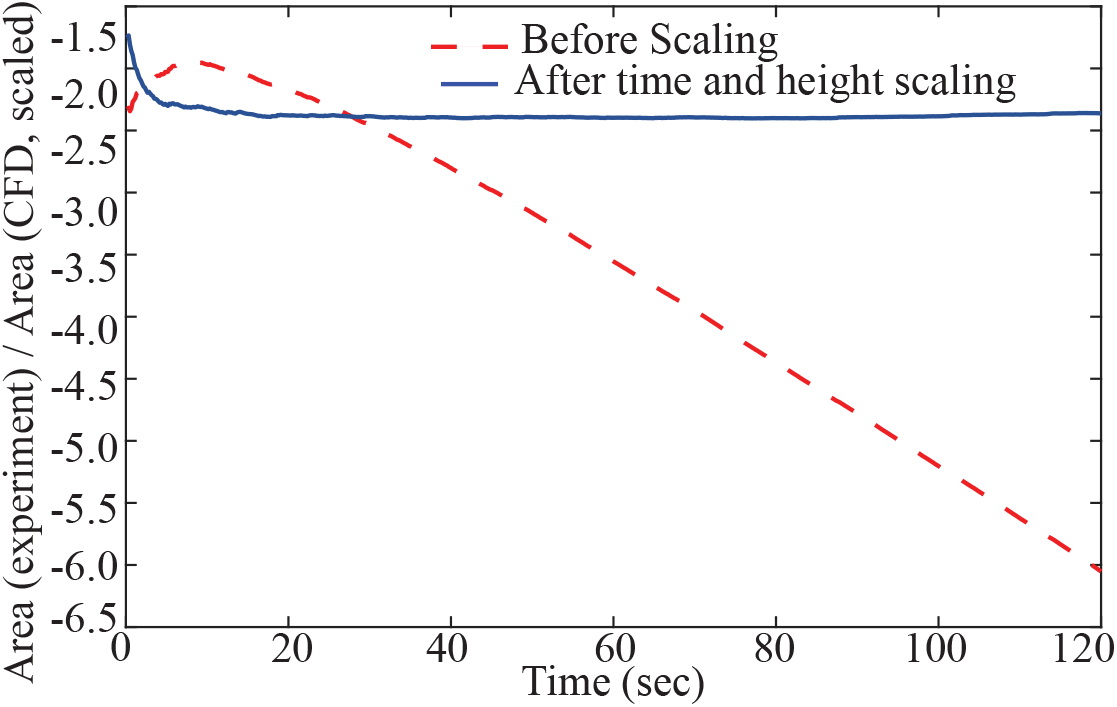
Scaling reconciliation of wetted-area growth between experiment and CFD: Wetted-area dynamics align between experiment and CFD after time and thickness scaling. Red dashed: experiment vs. raw CFD (before scaling). Blue solid: experiment vs. scaled CFD (after scaling). Post-scaling, the offset is nearly constant and the trends coincide, indicating matched advection-dominated growth.

After an initial transient, the residual flattens (blue curve), indicating consistent advection–dominated growth once thickness and time are reconciled; any remaining offset is attributable to early–time device effects (pump start–up/compliance, wetting/pinning and capillary entry on the resin surface, transient air evacuation) that are absent from the idealized 2D model.

### 4.3 On the use of a large inlet concentration in the RAD model

The reverse advection-diffusion (RAD) model matches CFD on both the advective and diffusion regimes, indicating that key transport physics are retained without full Navier–Stokes resolution; hence RAD is well-suited for rapid sweeps and design use. To theoretically model the diffusive component for the perfusion dynamics within the extracellular domain, we have imposed a finitely large value (*c*_in_ = 8.0 × 10^13^ in model units) for plasma concentration at time *t* = 0 as a pragmatic approximation [Crank, 1979] mimicking a persistent, high concentration fenestral ‘point’ source. A sufficiently large (but finite) value for initial inlet concentration creates a steep concentration gradient that effectively drives diffusion while maintaining the stability and robustness of the analytical solver. This approach aligns with standard practice in transport modeling, where high boundary concentrations are employed as Dirichlet boundary conditions to represent sustained sources, thus allowing for a realistic yet computationally feasible launch point for the complex intra-tumoral transport process without sacrificing physical relevance.

The present surrogate intentionally simplifies geometry and driving fields to isolate advection–diffusion obstruction physics: an idealized rectangle with uniformly sized, non-contacting circular fibers and a spatially simple inlet advection profile. Future work will incorporate histology-derived fiber statistics (size distribution, alignment, and contacts) and spatially varying ***ω***(*x, y*), informed by localized CFD or microfluidic measurements, thereby extending RAD from rapid screening toward tissue-specific predictions while preserving computational tractability.

## 5 Conclusion

We integrate three-phase, glycocalyx-patched (EHD) CFD with a histology-informed ECM domain to quantify how endothelial EHD alters plasma delivery at a fenestra and its downstream percolation. We then use a CFD-informed reverse advection–diffusion (RAD) model as a corroborative, geometry-aware surrogate for rapid exploration. The key findings are that across all 15 fenestra positions, adding EHD increases the inlet-side mean calibrated plasma intensity from 0.576 (Non-EHD) to 0.722 (EHD), representing a 25.34% gain. Downstream, both the ECM simulations and a microfluidic benchmark exhibit the same two-stage transport (advection-dominated entry followed by diffusion-driven spread), and the CFD-informed RAD reproduces these kinetics successfully.

In future work, we will extend the ECM-domain simulations to fully three-dimensional geometries, capturing transverse leakage paths and thickness-dependent effects that are absent in 2D. The reverse advection–diffusion model will be augmented with explicit electrohydrodynamic forcing (glycocalyx-patch charge and field–flow coupling) to incorporate EHD into the closures for effective advection and dispersion. Experimentally, we will develop refined organ-on-chip platforms with tunable wall charge and co-cultured endothelium–stroma to control EHD conditions in vitro, enabling direct quantification of perfusion trends under EHD versus Non-EHD and more rigorous validation of the modeling framework. Together, these advances will broaden biological realism and improve the predictive scope of EHD-aware tumor perfusion modeling.

## 6 Conflict of Interest Statement

The authors declare that the research was conducted in the absence of any commercial or financial relationships that could be construed as a potential conflict of interest.

## 7 Author Contributions

MMHA: geometry preparation, simulations, study design, data post-processing, data analysis, writing; MY: 3D printing, study design of RAD modeling, data analysis, writing; SM: running experiments; RAN: analysis, writing; JM: tumor scanning, contributions to writing; KR: tumor scanning, student supervision, contributions to writing; AP: experiment design, writing; SB: conceptualization, study design, funding acquisition, project administration, data analysis, writing.

## 8 Funding

This work was supported by the National Institutes of Health (NIH) Centers of Biomedical Research Excellence (COBRE) Award 2P20GM109024. Any opinions, findings, and conclusions or recommendations expressed here are, however, those of the authors and do not necessarily reflect views of the NIH.

## 9 Data Availability Statement

All essential information is contained in the article. Supplementary information (including anatomical geometries, simulation files, postprocessing spreadsheets, and programming codes) are available via the open access repository figshare, with doi: 10.6084/m9.figshare.30524669. The reader may also contact the corresponding author for any relevant data.

## Notes

https://figshare.com/s/bcbcae38be369f8ef98b

### Competing Interest Statement

The authors have declared no competing interest.

## References

Gullino, P. M. Tumor pathophysiology: the perfusion model. Antibiotics and Chemotherapy, 28:35–42, 1980.

Mostafa Sefidgar, Madjid Soltani, Kaamran Raahemifar, Hossein Bazmara, Seyed Mojtaba Mousavi Nayinian, and Majid Bazargan. Effect of tumor shape, size, and tissue transport properties on drug delivery to solid tumors. Journal of biological engineering, 8(1):12, 2014.

Vaupel, P.and Kallinowski, F. and Okunieff, P. Blood flow, oxygen consumption and tissue oxygenation of human tumors. Oxygen Transport to Tissue XII, pages 895–905, 1990.

Rakesh K Jain, John D Martin, and Triantafyllos Stylianopoulos. The role of mechanical forces in tumor growth and therapy. Annual review of biomedical engineering, 16(1):321–346, 2014.

Koumoutsakos, P., Pivkin, I., and Milde, F. The fluid mechanics of cancer and its therapy. Annual Review of Fluid Mechanics, 45(1):325–355, 2013.

Triantafyllos Stylianopoulos, Lance L Munn, and Rakesh K Jain. Reengineering the physical microenvironment of tumors to improve drug delivery and efficacy: from mathematical modeling to bench to bedside. Trends in cancer, 4 (4):292–319, 2018a.

Xianjie Jiang, Jie Wang, Xiangying Deng, Fang Xiong, Shanshan Zhang, Zhaojian Gong, Xiayu Li, Ke Cao, Hao Deng, Yi He, et al. The role of microenvironment in tumor angiogenesis. Journal of Experimental & Clinical Cancer Research, 39(1):204, 2020.

H. Maeda. Macromolecular therapeutics in cancer treatment: the EPR effect and beyond. Journal of Controlled Release, 164(2):138–144, 2012.

Stylianopoulos, T. and Jain, R. K. Combining two strategies to improve perfusion and drug delivery in solid tumors. Proceedings of the National Academy of Sciences, 110(46):18632–18637, 2013.

Gavhane, Y. N., Shete, A. S., Bhagat, A. K., Shinde, V. R., Bhong, K. K., Khairnar, G. A., and Yadav, A. V. Solid tumors: facts, challenges and solutions. Int J Pharm Sci Res, 2:1–12, 2011.

KM Taufiqur Rahman, Tanim Islam, Md Fahmid Islam, Roberto G Carbone, Nicholas C Butzin, and Md Khadem Ali. Targeting epigenetics in pulmonary arterial hypertension. In Targeting Epigenetics in Inflammatory Lung Diseases, pages 223–255. Springer, 2023.

Jain, R. K. Delivery of molecular medicine to solid tumors. Science, 271(5252):1079–1080, 1996.

Jain, R. K. The next frontier of molecular medicine: delivery of therapeutics. Nature Medicine, 4(6):655–657, 1998.

Letitia K Chim and Antonios G Mikos. Biomechanical forces in tissue engineered tumor models. Current Opinion in Biomedical Engineering, 6:42–50, 2018.

A. d’Esposito, P. W. Sweeney, M. Ali, M. Saleh, R. Ramasawmy, T. A. Roberts, G. Agliardi, A. Desjardins, M. F. Lythgoe, R. B. Pedley, et al. Computational fluid dynamics with imaging of cleared tissue and of in vivo perfusion predicts drug uptake and treatment responses in tumours. Nature Biomedical Engineering, 2(10):773–787, 2018.

M. Soltani and P. Chen. Numerical modeling of fluid flow in solid tumors. PloS One, 6(6):e20344, 2011.

Y. Zhang, L. Chen, J. Yang, J. B. Fleming, P. J. Chiao, C. D. Logsdon, and M. Li. Study human pancreatic cancer in mice: how close are they? Biochimica et Biophysica Acta (BBA) - Reviews on Cancer, 1835(1):110–118, 2013.

Sheldon Weinbaum, John M Tarbell, and Edward R Damiano. The structure and function of the endothelial glycocalyx layer. Annu. Rev. Biomed. Eng., 9(1):121–167, 2007.

Huilgo, R.R. and Phan-Thien, N. 7 - computational viscoelastic fluid dynamics. In Fluid Mechanics of Viscoelasticity, volume 6 of Rheology Series, pages 397–472. Elsevier, 1997. doi:10.1016/S0169-3107(97)80009-3. URL https://www.sciencedirect.com/science/article/pii/S0169310797800093.

PP Sumets, JE Cater, DS Long, and RJ Clarke. Electro-poroelastohydrodynamics of the endothelial glycocalyx layer. Journal of Fluid Mechanics, 838:284–319, 2018.

C Teodoro, J Arcos, O Bautista, and F Mendez. Electro-poroelastohydrodynamics of the endothelial glycocalyx layer and streaming potential in wavy-wall microvessels. Physical Review Fluids, 9(1):013101, 2024.

E. W. Merrill, G. C. Cokelet, A. Britten, and R. E. Wells Jr. Non-Newtonian rheology of human blood – effect of fibrinogen deduced by “subtraction”. Circulation Research, 13(1):48–55, 1963.

Baskurt, O. K. and Meiselman, H. J. Blood rheology and hemodynamics. Seminars in Thrombosis and Hemostasis, 29 (05):435–450, 2003.

Jonghwun Jung, Robert W Lyczkowski, Chandrakant B Panchal, and Ahmed Hassanein. Multiphase hemodynamic simulation of pulsatile flow in a coronary artery. Journal of Biomechanics, 39(11):2064–2073, 2006.

Mohammad Mehedi Hasan Akash, Nilotpal Chakraborty, Jiyan Mohammad, Katie Reindl, and Saikat Basu. Development of a multiphase perfusion model for biomimetic reduced-order dense tumors. Experimental and Computational Multiphase Flow, 5(3):319–329, 2023.

A. Rahmat. Numerical simulation of multiphase flows under electrohydrodynamics effects. PhD thesis, Sabanci University, 2017.

Hongyan Kang, Qiuhong Wu, Anqiang Sun, Xiao Liu, Yubo Fan, and Xiaoyan Deng. Cancer cell glycocalyx and its significance in cancer progression. International journal of molecular sciences, 19(9):2484, 2018.

J. W. Baish, T. Stylianopoulos, R. M. Lanning, W. S. Kamoun, D. Fukumura, L. L. Munn, and R. K. Jain. Scaling rules for diffusive drug delivery in tumor and normal tissues. Proceedings of the National Academy of Sciences, 108(5): 1799–1803, 2011.

M. Sugihara-Seki and B. M. Fu. Blood flow and permeability in microvessels. Fluid Dynamics Research, 37(1-2): 82–132, 2005.

Womersley, J. R. Oscillatory flow in arteries: the constrained elastic tube as a model of arterial flow and pulse transmission. Physics in Medicine & Biology, 2(2):178, 1957.

Attinger, E. O. Elements of Theoretical Hydrodynamics, pages 15–76. New York, McGraw-Hill, 1964.

Srivastava, V. P. A theoretical model for blood flow in small vessels. Applications and Applied Mathematics: An International Journal (AAM), 2(1):5, 2007.

M. Sefidgar, M. Soltani, K. Raahemifar, and H. Bazmara. Effect of fluid friction on interstitial fluid flow coupled with blood flow through solid tumor microvascular network. Computational and Mathematical Methods in Medicine, 2015, 2015.

Mohammad Mehedi Hasan Akash, Mohammad Yeasin, Pei Ran, Anna-Blessing Merife, Anupam Pandey, Pranav Soman, and Saikat Basu. Modeling transport in physiologically realistic tumor microenvironment. In APS Division of Fluid Dynamics Meeting Abstracts, pages L04–001, 2024.

Akash, M. M. H., Chakraborty, N., and Basu, S. A multiphase tracking of perfusion through in silico dense tumor domain. In APS Division of Fluid Dynamics Meeting Abstracts, pages N01–061, 2021.

P. Nithiarasu. Biofluid Dynamics, chapter 2, pages 20–21. Electronic book: Download source, accessed 12-October-2022, 2022.

Randal O Dull and Robert G Hahn. The glycocalyx as a permeability barrier: basic science and clinical evidence. Critical Care, 26(1):273, 2022.

Sheldon Weinbaum, Limary M Cancel, Bingmei M Fu, and John M Tarbell. The glycocalyx and its role in vascular physiology and vascular related diseases. Cardiovascular engineering and technology, 12(1):37–71, 2021.

Eekhoff, J. D. and Lake, S. P. Three-dimensional computation of fibre orientation, diameter and branching in segmented image stacks of fibrous networks. Journal of the Royal Society Interface, 17(169):20200371, 2020.

Gidaspow, D. Multiphase Flow and Fluidization: Continuum and Kinetic Theory Descriptions. Academic Press, 1994.

Anderson, T. B. and Jackson, R. Fluid mechanical description of fluidized beds. Industrial & Engineering Chemistry Fundamentals, 6(4):527–539, 1967.

Schwartz, R. S. and Conley, C. L. Blood. Encyclopedia Britannica, 2020. URL https://www.britannica.com/science/blood-biochemistry.

Lev Davidovich Landau, John Stewart Bell, MJ Kearsley, LP Pitaevskii, EM Lifshitz, and JB Sykes. Electrodynamics of continuous media, volume 8. elsevier, 2013.

DA1435033 Saville. Electrohydrodynamics: the taylor-melcher leaky dielectric model. Annual review of fluid mechanics, 29(1):27–64, 1997.

Antonio Castellanos, Antonio Ramos, Antonio Gonzalez, Nicolas G Green, and Hywel Morgan. Electrohydrodynamics and dielectrophoresis in microsystems: scaling laws. Journal of Physics D: Applied Physics, 36(20):2584, 2003.

Todd M Squires and Martin Z Bazant. Breaking symmetries in induced-charge electro-osmosis and electrophoresis. Journal of Fluid Mechanics, 560:65–101, 2006.

H Pauly and HP Schwan. Dielectric properties and ion mobility in erythrocytes. Biophysical journal, 6(5):621–639, 1966.

Alexander Zhbanov and Sung Yang. Effects of aggregation on blood sedimentation and conductivity. PloS one, 10(6): e0129337, 2015.

Jun Yang, Ying Huang, Xujing Wang, Xiao-Bo Wang, Frederick F Becker, and Peter RC Gascoyne. Dielectric properties of human leukocyte subpopulations determined by electrorotation as a cell separation criterion. Biophysical journal, 76(6):3307–3314, 1999.

J. Jung and A. Hassanein. Three-phase CFD analytical modeling of blood flow. Medical Engineering & Physics, 30(1): 91–103, 2008.

Jung, J., Hassanein, A., and Lyczkowski, R. W. Hemodynamic computation using multiphase flow dynamics in a right coronary artery. Annals of Biomedical Engineering, 34(3):393, 2006.

Z. Naumann and L. Schiller. A drag coefficient correlation. Z. Ver. Deutsch. Ing, 77(318):e323, 1935.

Zhao, G., Wu, J., Xu, S., Collins, M., Long, Q., Jian König, Carola S. and Jiang, Yuping and Wang, and Padhani, AR. Numerical simulation of blood flow and interstitial fluid pressure in solid tumor microcirculation based on tumor-induced angiogenesis. Acta Mechanica Sinica, 23(5):477–483, 2007.

Afolabi, E. A. and Lee, J. An Eulerian-Eulerian CFD simulation of air-water flow in a pipe separator. The Journal of Computational Multiphase Flows, 6(2):133–149, 2014.

Wu, M., Frieboes, H. B., McDougall, S. R., Chaplain, M. AJ., Cristini, V., and Lowengrub, J. The effect of interstitial pressure on tumor growth: coupling with the blood and lymphatic vascular systems. Journal of Theoretical Biology, 320:131–151, 2013.

Sven, K. and Josipa, F. Interstitial hydrostatic pressure: a manual for students. Advances in Physiology Education, 31 (1):116–117, 2007.

T. Stylianopoulos, L. L. Munn, and R. K. Jain. Reengineering the physical microenvironment of tumors to improve drug delivery and efficacy: from mathematical modeling to bench to bedside. Trends in Cancer, 4(4):292–319, 2018b.

Heldin, C., Rubin, K., Pietras, K., and Östman, A. High interstitial fluid pressure – an obstacle in cancer therapy. Nature Reviews Cancer, 4(10):806–813, 2004.

Porterfield, W. W. and Kruse, W. Loschmidt and the discovery of the small. Journal of Chemical Education, 72(10):870, 1995.

R. Andersson, W. G. Bouwman, S. Luding, and I. M. De Schepper. Stress, strain, and bulk microstructure in a cohesive powder. Physical Review E, 77(5):051303, 2008.

V. Nier, S. Jain, C. T. Lim, S. Ishihara, B. Ladoux, and P. Marcq. Inference of internal stress in a cell monolayer. Biophysical Journal, 110(7):1625–1635, 2016.

C. P. Spatarelu, H. Zhang, D. T. Nguyen, X. Han, R. Liu, Q. Guo, J. Notbohm, J. Fan, L. Liu, and Z. Chen. Biomechanics of collective cell migration in cancer progression: experimental and computational methods. ACS Biomaterials Science & Engineering, 5(8):3766–3787, 2019.

Mohammad Yeasin. Integrative theoretical-numerical modeling of plasma transport and perfusion in solid tumors. 2025.

Mats Leeman, Jaeyeong Choi, Sebastian Hansson, Matilda Ulmius Storm, and Lars Nilsson. Proteins and antibodies in serum, plasma, and whole blood—size characterization using asymmetrical flow field-flow fractionation (af4). Analytical and bioanalytical chemistry, 410(20):4867–4873, 2018.

Noriaki Muraoka, Hidemasa Uematsu, Hirohiko Kimura, Yoshiaki Imamura, Yasuhiro Fujiwara, Makoto Murakami, Akio Yamaguchi, and Harumi Itoh. Apparent diffusion coefficient in pancreatic cancer: characterization and histopathological correlations. Journal of Magnetic Resonance Imaging, 27(6):1302–1308, 2008.

Jacob Cohen. Statistical power analysis for the behavioral sciences. routledge, 2013.

John Crank. The mathematics of diffusion. Oxford university press, 1979.

Muhammad Sahimi. Flow and transport in porous media and fractured rock: from classical methods to modern approaches. John Wiley & Sons, 2011.

Ralf Metzler and Joseph Klafter. The random walk’s guide to anomalous diffusion: a fractional dynamics approach. Physics reports, 339(1):1–77, 2000.

Brian Berkowitz, Harvey Scher, and Stephen E Silliman. Anomalous transport in laboratory-scale, heterogeneous porous media. Water Resources Research, 36(1):149–158, 2000.

Yaniv Edery, Alberto Guadagnini, Harvey Scher, and Brian Berkowitz. Origins of anomalous transport in heterogeneous media: Structural and dynamic controls. Water Resources Research, 50(2):1490–1505, 2014.

